# H3K4 methylation-promoted transcriptional memory ensures faithful zygotic genome activation and embryonic development

**DOI:** 10.1101/2025.01.20.633863

**Authors:** Meghana S. Oak, Marco Stock, Matthias Mezes, Tobias Straub, Antony M. Hynes-Allen, Jelle van den Ameele, Ignasi Forne, Andreas Ettinger, Axel Imhof, Antonio Scialdone, Eva Hörmanseder

## Abstract

In the life of a vertebrate embryo, gene expression is initiated for the first time at zygotic genome activation (ZGA). Maternally expressed transcription factors present in the embryo are essential for this process. However, it is unknown if active chromatin modifications established in the gamete are propagated in the embryo as an epigenetic memory to support ZGA and embryonic development. Here, we provide evidence that in *Xenopus laevis*, H3K4 methylation provides an epigenetic memory of active chromatin states. We show that this is required for faithful zygotic genome activation and successful embryonic development. Chromatin configurations of promoters displaying high H3K4me3 intensity and breadth, alongside DNA hypomethylation and increased GC content, are propagated from the gametes to the embryo across multiple cell divisions and a transcriptionally quiescent phase in early development. We show that this transmission of H3K4 methylation is essential for precise zygotic genome activation and expression of key pioneer ZGA transcription factors Pou5f3.2 and Sox3. Finally, we demonstrate that the H3K4 methyltransferases Kmt2b and Cxxc1 ensure transcription-independent propagation of H3K4me3 and proper zygotic gene expression. In summary, our study establishes the role of H3K4 methylation in maintaining memory of active chromatin states in *Xenopus* embryos and reveals its importance for successful embryonic development.

## Introduction

Cell fates are established through the concerted activity of developmental cues, transcription factors (TFs) and a multitude of epigenetic factors that either permit or repress specific transcriptional programs. In vertebrates, early development of the zygote is supported by maternal transcripts and TFs, until zygotic genome activation (ZGA), when the first gene expression patterns of the embryo are established to ultimately define cell fate. The extent to which the establishment of these gene expression patterns depends on TFs or epigenetic mechanisms, or both, is not fully understood. Similarly, it is unclear whether the landscape of parental chromatin contributes to embryonic development via the epigenetic memory of chromatin states.

Epigenetic memory refers to the maintenance of chromatin states across cell divisions independently of the developmental signal that induced the state^1^. To date, most studies of epigenetic inheritance have confirmed the transmission of repressive histone modifications around transcriptionally inactive regions of chromatin. However, the role of active histone modifications—those traditionally associated with actively transcribed genes—in epigenetic memory is under debate^2^. It is generally considered that the maintenance of active chromatin states is dependent on transcription factor-coupled gene expression^3^. Propagation of active histone marks is thought to be dependent on transcription factor binding and continuous gene transcription, and not via direct transmission using autonomously operating read-and-write mechanisms. Therefore, the field does not unequivocally consider the maintenance of active histone marks as an epigenetic mechanism. Moreover, the close association of active chromatin states with transcription creates a great challenge for experiments aiming to disentangle whether active histone marks represent a cause or a consequence of transcription, and whether they might embody epigenetic mechanisms^4,5^.

Previous studies in fast-developing embryos such as *Xenopus laevis* and zebrafish point to differing models. Embryos of these species are of particular interest for studying maintenance of active epigenetic marks, as they undergo a prolonged transcription-free window of at least 8 rapid cell divisions from fertilization to zygotic genome activation (ZGA). This poses a challenging environment for retention of chromatin modifications^6^. In zebrafish, the active chromatin mark H3K4me3 has been found to be present before ZGA at developmental gene promoters, and placeholder mechanisms driven by other chromatin modifications, namely H2A.Z and DNA methylation, were shown to set the stage for the first transcriptional program^7,8^. In *Xenopus*, technical limitations hamper the design of experiments analyzing histone marks at these early developmental stages. Thus, the presence of H3K4me3 on this highly dynamic early embryonic chromatin is questioned to date^9^. Instead, it has been suggested that H3K4me3 is rapidly acquired *de novo* around ZGA in blastula embryos^9–11^. This model would exclude a direct role of H3K4me3 in transmitting active chromatin states from gametes to embryos, as well as overlook the contribution of such a mechanism for successful embryonic development.

In contrast and similar to the abovementioned zebrafish findings, our previous experiments revealed that H3K4me3 decorates specific regions of the genome in *Xenopus* sperm nuclei and in blastula embryos prior to ZGA^12^. This opens the possibility that other chromatin marks could also be present on the highly dynamic chromatin of pre-ZGA embryos. In addition, cell fate reprogramming studies performing somatic cell nuclear transfer in *Xenopus* embryos suggest that H3K4me3 may play a role in transmission of active transcriptional states, independently of continuous transcription^13^. Specifically, H3K4 demethylation of somatic donor cell nuclei rendered them permissive to reprogramming, and transcriptional memory of active gene expression states was lost^13^. These experiments excitingly suggested that nuclear transfer embryos retain information encoded by H3K4me3 via epigenetic memory. Moreover, it raised the question whether this mechanism also propagates information from the gametes to the fertilized embryo during physiological development. Together, these observations support the possibility of the involvement of H3K4me3 in transmission of active transcriptional states between fertilization and ZGA, but concrete evidence is currently missing. Furthermore, it has not been tested if such maintenance of active chromatin states via H3K4 methylation would be important for successful embryonic development.

Here, we made use of the transcriptionally silent pre-ZGA developmental period in *Xenopus laevis* embryos to investigate the transmission of active histone modifications independently of transcription, and its role in embryonic development. To achieve this, we profiled histone modifications on chromatin in pre-ZGA embryos using mass spectrometry. We detected H3K4me3 together with a plethora of other histone modifications associated with active chromatin on pre-ZGA chromatin, despite multiple rapid cell divisions in the absence of transcription. We then traced the H3K4me3 mark on a large group of genes that are transcribed in the parental gametes, as well as the ZGA embryo, separated by transcriptional dormancy in early cleavage - stage embryos. We show that H3K4me3 peaks are maintained around these genes across pre-ZGA development. The promoters of these genes are furthermore characterized by high CpG density and DNA accessibility, as well as DNA hypo-methylation, when compared with other genes. Knockdown of the transcription-independent H3K4 methyltransferases Kmt2b or Cxxc1 results in reduction of H3K4me3 levels, improper gene expression of zygotic genes and embryonic defects. Importantly, H3K4 demethylation specifically during pre-ZGA cleavages shows defects of zygotic genome activation that are comparable to those caused by prolonged H3K4 demethylation extending across ZGA. This suggests that the transcription-independent propagation of H3K4me3 during early embryonic cell cycles is essential for faithful ZGA. Together, this study demonstrates a role of H3K4 methylation as an epigenetic memory factor important for embryonic development.

## Results

### Histone modifications associated with active chromatin are present in early embryos despite rapid cell divisions and absence of transcription

After fertilization, the *Xenopus laevis* zygote divides rapidly at least 8 times before gene transcription is fully activated at ZGA. We tested first whether chromatin marks associated with transcriptionally active chromatin states are maintained on such dynamic, fast replicating chromatin in the absence of gene transcription.

To this end, we established a new nuclear extraction method as a prerequisite to our chromatin profiling experiments, which allowed us to overcome technical difficulties prevalent in the field arising from interference of high proportions of yolk protein in early *Xenopus laevis* embryos. Using this approach, we could profile the chromatin using mass spectrometry analysis of histone tail post-translational modifications for the first time in pre-ZGA vertebrate embryos (see methods section and **Fig.1a**). We detected a wide range of active and repressive histone modifications on globally transcriptionally silent chromatin of pre-ZGA embryos (256-cell stage). These include the active marks H3K4me1/2/3, H3K9ac and H3K27ac, but not H3K36me3, and the repressive marks H3K9me3 and H3K27me3 (**Fig.1b).** As expected, we also observed these histone marks on transcriptionally active chromatin of mid-ZGA embryos (4,000-cell stage; **Fig.1b**). Importantly, all three methylation states of the H3K4 residue can be detected on chromatin of pre- and mid-ZGA embryos, with an increase in the abundance of each modification as ZGA progresses (**Fig.1c**). As expected, the replication-associated histone modification H4K20me1 is highly abundant in both pre-ZGA and mid-ZGA embryos, during which embryos undergo extensive DNA replication and duplicate their genome every 30 minutes (**Fig.1b**). Together, these results reveal for the first time that the highly dynamic chromatin of rapidly dividing pre-ZGA embryos is decorated by both repressive and active chromatin marks.

**Figure 1.**
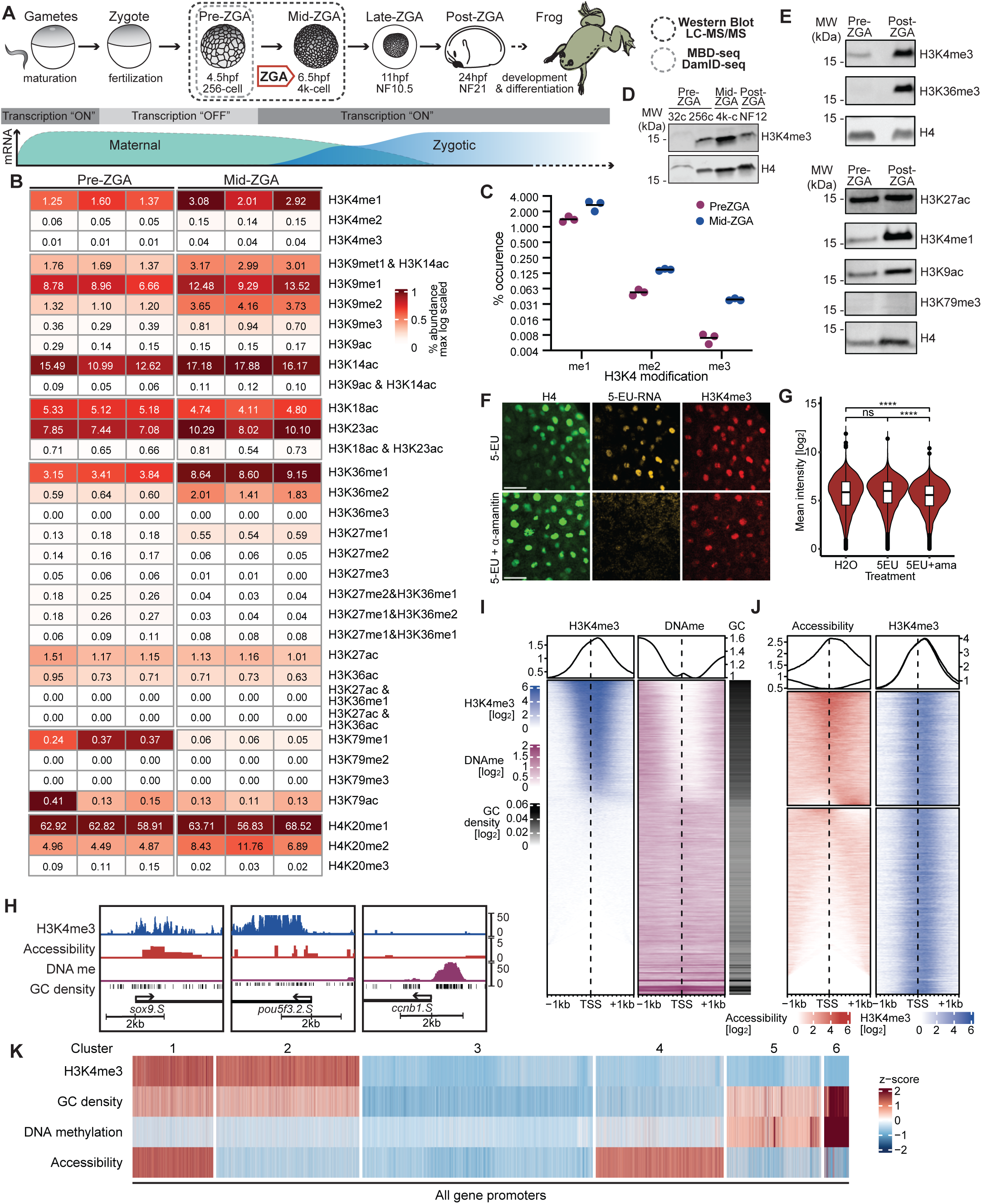
Profiling histone modifications and epigenetic factors present on pre-ZGA embryo chromatin. (A) Schematic demonstrating nuclear isolation on pre-ZGA embryos. (B) Heatmap showing the percent abundance of histone modifications on each respective peptide obtained by PTM analysis of LC-MS/MS. Numbers displayed are percent abundance of the modification, colors are max log scaled per peptide. (C) Percent abundance of modifications of H3K4 peptide at pre-ZGA and ZGA stages. (D) Time-series Western blot showing occurrence of H3K4me in embryos. (E) Western blots of histone modifications in pre-ZGA and post-ZGA embryos using nuclear isolation. (F) Representative images of H3K4me3 levels in wildtype and transcription-inhibited mid-ZGA embryos. Nascent transcripts are labeled using 5-EU. Nuclei are segmented using H4 (green), and H3K4me3 (red) and EU-RNA (yellow) values are measured. Scale bars: 50µm. (G) Quantification of immunofluorescence based on H3K4me3 signal in segmented nuclei. Statistical test: one-sided Wilcoxon rank-sum test; p-values: (****)<=0.0001, (***)<=0.001, (**) <= 0.01, (*)<=0.05; n.s. are p-values > 0.05. (H) IGV tracks displaying H3K4me3 ChIP-seq, CATaDa, and MBD-seq data for pre-ZGA embryos with CpG regions at representative promoters (TSS+/−1kb). (I) Heatmap showing H3K4me3 and DNA methylation at all gene promoters (TSS+/−1kb) in pre-ZGA embryos (4.5hpf), ordered by H3K4me3 enrichment. Promoter CpG density also shown (right). (J) Heatmap showing accessibility and H3K4me3 enrichment at all gene promoters (TSS+/−1kb) in pre-ZGA embryos (4.5hpf), ordered by accessibility enrichment and clustered using k-means clustering (k=2). (K) Heatmap showing z-scored H3K4me3 enrichment, CG density, DNA methylation enrichment and accessibility enrichment across promoters (TSS+/−1kb) in pre-ZGA embryos and clustered by k-means clustering (k=6; Hartigan-Wong algorithm).

We then confirmed these results by Western Blotting of isolated embryo nuclei. Importantly, we detected H3K4me3 modified chromatin in pre-ZGA embryos from as early as the 32-cell stage in development (**Fig.1d**). At the 256-cell stage, the active histone marks H3K4me3 as well as H3K4me1, H3K9ac and H3K27ac were detectable on mostly transcriptionally silent chromatin at levels comparable to those observed on transcriptionally active chromatin of a post-ZGA stage embryo (**Fig.1e**). In contrast, we did not detect the active histone marks H3K36me3 and H3K79me3 in pre-ZGA embryo nuclei, but only in transcribing post-ZGA nuclei. These results support that active histone marks H3K4me1/3, H3K9ac and H3K27ac are present on chromatin prior to ZGA.

In *Xenopus*, as well as in drosophila, zebrafish and mouse, a small number of microRNAs and β-catenin targets are transcribed before the major wave of ZGA—as early as the 8-cell stage and 128-cell embryos, respectively^14–18^. We hypothesized that such underlying early transcription may reinforce H3K4 promoter methylation through transcription-coupled mechanisms^4,19^. Therefore, we tested next whether inhibition of transcription by injecting 1-cell embryos with a high dose of the RNA Polymerase II and III inhibitor ⍺-amanitin results in acute depletion of H3K4me3 in embryos during ZGA (**Fig.S1a**). We co-injected the uridine analog 5-EU, which is incorporated into nascent RNAs^20^ to reveal effective inhibition of transcription (**Fig. S1b**). We observed that both H3K4me3 and H4 are detected in transcription-inhibited, 5-EU negative embryos and transcriptionally active, 5-EU positive embryos at developmental stages corresponding to mid-ZGA **(Fig.1f, Fig.1g**). We note that H3K4me3 levels are moderately reduced by 26% in ⍺-amanitin-injected embryos, indicating that transcription is indeed required for a proportion of this modification **(Fig.1g)**. Together, this suggests that the great majority of H3K4me3 marks on chromatin of mid-ZGA embryos are independent of RNA Pol II and III-mediated transcription.

These results demonstrate that the dynamic chromatin of rapidly dividing, and transcriptionally quiescent *Xenopus* embryos is decorated by H3K4me3, together with several other active and repressive histone modifications. This also indicates that during developmental stages preceding zygotic genome activation, chromatin states are established and maintained by mechanisms which operate independently of feedback from ongoing gene transcription.

### H3K4me3-marked promoters in pre-ZGA embryos are in distinct chromatin configurations

Having established that pre-ZGA chromatin is extensively marked with active histone modifications, we then addressed the genomic context of pre-ZGA chromatin in relationship to H3K4me3. In transcriptionally active cells, H3K4me3 is enriched around promoter regions of expressed genes and has been associated with DNA hypomethylation, high CpG density and accessible chromatin^21–25^. We tested if these associations are also present around the promoters of transcriptionally inactive genes in embryos prior to zygotic genome activation.

To this end, we profiled H3K4me3, DNA methylation and accessibility of chromatin in pre-ZGA embryos respectively by ChIP-seq, MBD-seq and CATaDa (Chromatin Accessibility profiling using Targeted Dam-ID) (**Fig.1a, Fig.1h)**^26,27^ and calculated CpG density and occurrence of H3K4me3, DNA methylation and accessibility of the *Xenopus* genome at promoter regions **(Fig.S1c).** We observed that H3K4me3 coincides with low levels of DNA methylation and high CpG densities around promoter regions of transcriptionally inactive genes in pre-ZGA embryos (**Fig.1i, k** clusters 1 and 2). 38% of the H3K4me3-marked promoters were found in an accessible configuration (**Fig.1j),** DNA hypomethylated and CpG-rich (**Fig.1k** cluster 1) and were thus in a similar chromatin configuration as transcriptionally active genes. The remaining H3K4me3 marked promoters were less accessible, but still DNA hypomethylated and with high CpG density (**Fig.1j, Fig.1k** cluster 2). Promoters that lacked H3K4me3 were in one out of three different configurations: DNA methylated, CpG-rich with low accessibility, suggesting a repressive chromatin state (**Fig.1k** clusters 5 and 6), DNA hypomethylated with low CpG density and accessible (**Fig.1k** cluster 4), or DNA hypomethylated with low CpG density and low accessibility (**Fig.1k** cluster 3).

Together, this shows that in *Xenopus* embryos at stages that precede ZGA, H3K4me3-marked promoters of transcriptionally silent genes are in distinct chromatin states regarding CpG content, DNA methylation and chromatin accessibility.

### Genes with high H3K4me3 intensity and breadth around their promoters maintain this state in early embryos

We then investigated the origins and fate of H3K4me3-marked promoters identified in pre-ZGA embryos. This was done by tracing the genomic landscape of H3K4me3 from the last transcriptionally active stages in the gametes to the next transcriptionally active stage in the embryo at ZGA. This developmental window spans gamete maturation, fertilization, and a transcription-free period during early cleavages of the embryo until the onset of zygotic transcription (**Fig.2a**). For this, we utilized published H3K4me3 chromatin immunoprecipitation (ChIP) sequencing datasets for spermatid, sperm, pre- and post-ZGA embryonic stages^12,28,29^. Low chromatin-to-yolk ratios in the egg prevented us from profiling H3K4me3 in the maternal genome, hence we can only deduct the maintenance of the mark from the paternal genomes.

**Figure 2.**
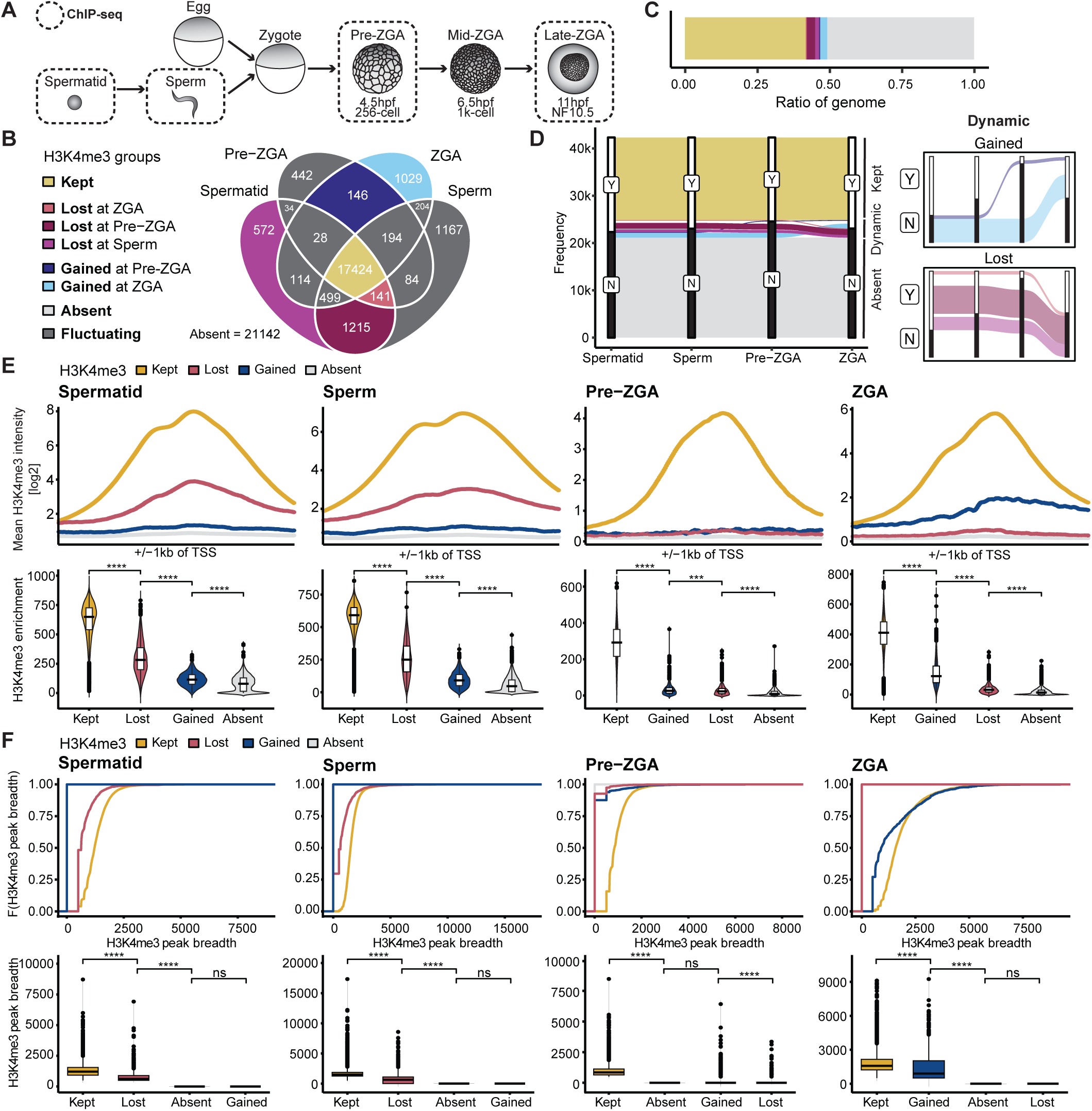
H3K4me3 peak intensity closely correlates with the duration of H3K4me3 maintenance. (A) Illustration highlighting stages of interest for H3K4me3 ChIP-seq. (B) Venn diagram denoting overlap of H3K4me3 promoter peaks across timepoints. Colors are based on the dynamics of H3K4me3 across the stages: Yellow: genes that “maintain” H3K4me3 (“KEPT”), blue: genes that “gain” H3K4me3 (“GAINED”), purple: genes that “lose” H3K4me3 peaks across timepoints (“LOST”). Promoters with fluctuating H3K4me3 dynamics are excluded from further analysis and marked in dark grey. (C) Ratio of genes that fall into each group of H3K4me3 dynamics. (D) Alluvial plots representing H3K4me3 dynamics of each group. (E) (Top) H3K4me3 around TSS (+/−1kb) for H3K4me3 dynamics groups at each timepoint - spermatid, sperm, pre-ZGA, post-ZGA respectively. (Bottom) H3K4me3 promoter enrichment in each group for each timepoint. Statistical test: one-sided Wilcoxon rank-sum test; p-values: (****)<=0.0001, (***)<=0.001, (**) <= 0.01, (*)<=0.05; n.s. are p-values > 0.05. (F) Cumulative distributions (ECDF plots) (top) and distributions (bottom) of H3K4me3 peak breadth at promoters in each H3K4me3 dynamics group (bp) at each time point - spermatid, sperm, pre-ZGA, post-ZGA respectively.

We found that the called H3K4me3 peaks predominantly associated with gene promoter regions in the gametes and early embryonic samples (**Fig.S2a**). Importantly, 75% of all identified H3K4me3 promoter peaks overlapped across all four time points and made up 39% of all gene promoters **(“**KEPT” group**; Fig.2b, Fig.2c)**, raising the possibility that H3K4me3 is established in the spermatid stage and is then maintained at the promoters of these genes until ZGA. In contrast, for 48% of all genes, H3K4me3 peaks were not detected at their promoters at any of the four time points (“ABSENT” group, **Fig.2c, Fig.2d)**. Only the remaining 13% of all genes showed dynamic behavior in regard to H3K4me3 at their promoters across the analyzed timepoints (**Fig.2b, Fig.2c, Fig.2d, Fig.S2f)** and were further classified based on their patterns. Genes within the “LOST” group had an H3K4me3 peak at the spermatid stage, which is lost during one of the subsequent developmental stages **(Fig.2b, Fig.2d, Fig.S2b)**. Conversely, genes within the “GAINED” group lacked an H3K4me3 promoter peak in the spermatid or sperm stage but gained it predominantly at ZGA stages, with only a small subset gaining it already at pre-ZGA **(Fig.2b, Fig.2d, Fig.S2b)**. These results suggest that the majority of H3K4me3-marked promoters identified in pre-ZGA stage embryos are already in this state in the male gametes and maintain this mark in ZGA stage embryos.

We then evaluated H3K4me3 peak characteristics of our gene groups representing differing H3K4me3 dynamics across development. We find that the KEPT group shows significantly higher H3K4me3 peak intensities and peak breadths around the promoter regions than all other groups across all analyzed timepoints **(Fig.2e, Fig.2f)**. Furthermore, we see that the relative H3K4me3 promoter enrichment and breadth in the spermatid indicate how long the mark remains detectable during the subsequent developmental stages. Specifically, the promoters of the KEPT group of genes, which maintain H3K4me3 marks for the longest duration of time, show highest H3K4me3 enrichment and breadth scores in spermatid **(Fig.S2c)**, followed by the promoters that show a loss at ZGA, at pre-ZGA and at sperm stage, which show progressively lower scores. At the ZGA stage, the promoters that gain H3K4me3 peaks at the pre-ZGA stage show greater breadth and intensity of H3K4me3 enrichment compared to peaks that are gained later at the ZGA stage (**Fig.S2d**). H3K4me3 extending into the coding region has been described for actively transcribed genes in somatic cells^30^, and we also observed asymmetrical extension of peak breadth into the gene body for GAINED genes during onset of ZGA **(Fig.2e).** In summary, these analyses reveal that H3K4me3 peak intensities and peak breadth closely correlate with the duration of H3K4me3 maintenance in early development.

Finally, we addressed what kind of genes fall into these identified groups with specific H3K4me3 dynamics. Gene ontology analyses revealed an enrichment of terms associated with housekeeping functions mainly in the KEPT group, whereas terms related to embryo development, gastrulation and pattern specification were highly enriched in the GAINED group, and to a lesser extent in the KEPT group (**Fig.S2e)**. The LOST group did not show significant enrichment for any biological processes. Thus, in the embryo, propagation of H3K4me3 at promoter regions may be important for genes with housekeeping functions, whereas a gain in H3K4me3 at the ZGA stage may be important for genes with development and differentiation functions. This aligns with known gene expression patterns during early *Xenopus* development.

Together, these analyses reveal that H3K4me3 domains around promoters of genes that display high intensity and breadth are maintained from the last transcriptionally active stages in the gamete, across several, largely transcriptionally inactive cell divisions, until the next transcriptionally active stage ZGA stage in the embryo.

### Genes transcribed in gametes and during ZGA preferentially maintain H3K4me3 during the interjacent transcriptionally silent phase

Since our analyses indicate H3K4me3 maintenance around genes from gametes to embryos across a developmental time window where they are transcriptionally silent, we addressed next whether this maintenance could contribute to inheriting memory of past active transcriptional states. For that to be the case, we hypothesized that genes that maintain H3K4me3 around their promoters in gametes and in embryos were once actively transcribed in the gametes and are then transcribed again during ZGA.

We thus asked whether we could identify genes that were expressed in the gametes and expressed again during ZGA in embryos. For this, we made use of published datasets of nascent RNA-seq spanning pre- and mid-ZGA stages of embryonic development (5-9hpf) as well as total RNA sequencing datasets of the gamete precursors, egg and spermatid, to characterize specific gene expression groups based on their transcriptional dynamics^28,29,31^. Indeed, we refer to genes whose transcripts are detected in either the maternal or paternal gamete, as well as in the embryo during the above ZGA timepoints as *“Gamete and Zygotically expressed genes” (GZ genes,* **Fig.3a, Fig.3b, Fig.3c**). In addition, we identified genes exclusively expressed in the maternal and paternal gametes (*“Gamete-Specific”, GS* genes) as well as genes that are expressed during ZGA and are not detected in either gamete (*“Zygote-Specific”, ZS* genes; **Fig.3a, Fig.3b, Fig.3c**). The group of genes whose transcripts are undetectable in all the above time points are denoted as *“Not-Detected” (ND)* genes; these genes may be expressed at later time points in development. Gene ontology (GO) analyses revealed that the term “embryo development” is associated with both GZ and ZS genes. Additionally, in agreement with our analysis of H3K4me3 dynamics, terms related to housekeeping functions and mRNA processing are primarily linked to GZ genes, while terms associated with gastrulation, morphogenesis and symmetry formation are specifically enriched in ZS genes **(Fig.S3a-d).** Together, this suggests that genes in the GZ group may propagate their active transcriptional state from the gametes to the ZGA embryo and have potential housekeeping or developmental functions.

**Figure 3.**
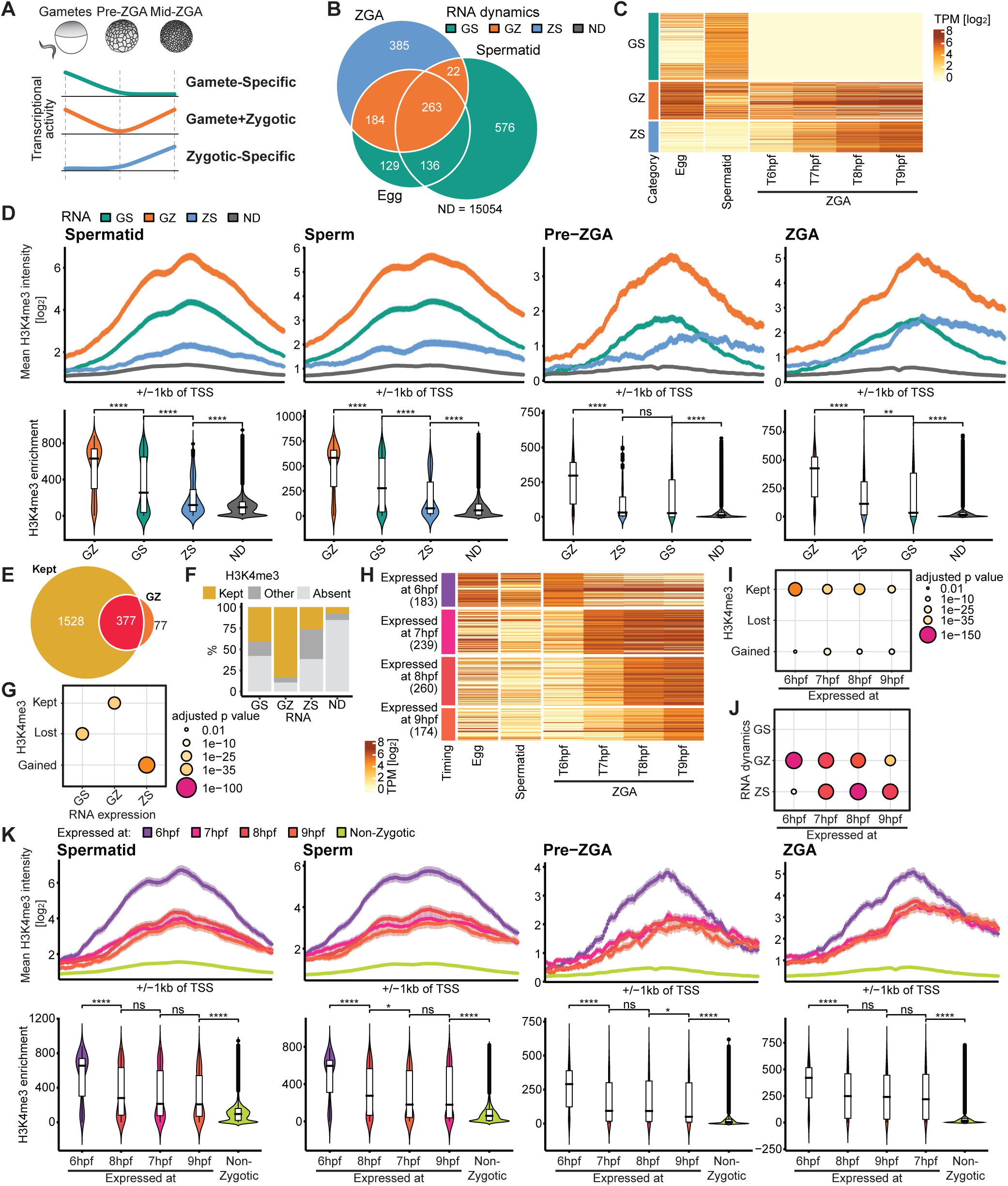
H3K4me3 is maintained independently of transcription between two transcriptionally active time points. (A) Schematic representing transcriptional activity in gametes, pre-ZGA and ZGA embryos in each RNA group. *Gamete-Specific (GS)*: genes that are only expressed in the gametes, *Gamete-Zygotic (GZ)*: genes that are expressed in the gametes and zygotically, *Zygotic-Specific (ZS)*: genes are expressed only zygotically. (B) Venn diagram denoting overlap of genes detected in the egg, spermatid (total RNA) and at ZGA stage (nascent). Colors are based on the dynamics of RNA expression across stages. (C) Heatmap of RNA expression (log scaled TPM) of the different RNA dynamics groups in the egg, spermatid and different timepoints during ZGA (6-9hpf). (D) (Top) H3K4me3 around TSS (+/−1kb) for RNA dynamics groups at each timepoint - spermatid, sperm, pre-ZGA, post-ZGA respectively. (Bottom) H3K4me3 promoter enrichment in each group for each timepoint. Statistical test: one-sided Wilcoxon rank-sum test; p-values: (****)<=0.0001, (***)<=0.001, (**) <= 0.01, (*)<=0.05; n.s. are p-values > 0.05. (E) Overlap of KEPT H3K4me3 dynamics group and the GZ RNA dynamics group. (F) Composition of each RNA dynamics group with respect to the H3K4me3 dynamics groups. LOST and GAINED are summarized as OTHER. (G) Results of statistical tests for association between the H3K4me3 dynamics groups and RNA dynamics groups. P-values are calculated with one-sided Fisher’s Exact Test and FDR adjusted. (H) Heatmap of expression of genes in egg, spermatid, and different timepoints during ZGA (6-9hpf) grouped by their first detected expression timepoint during ZGA. (I-J) Results of statistical tests for association between (I) H3K4me3 dynamics groups and the first detected expression timepoint during ZGA (6-9hpf) and (J) RNA dynamics groups and the first detected expression timepoint during ZGA (6-9hpf). P-values are calculated with one-sided Fisher’s Exact Test and FDR adjusted. (K) (Top) H3K4me3 around TSS (+/−1kb) for ZGA timing groups at each timepoint - spermatid, sperm, pre-ZGA, post-ZGA respectively. (Bottom) H3K4me3 promoter enrichment in each group for each timepoint.

To address whether the promoters of the GZ group of genes are continuously marked by H3K4me3, we calculated and compared the H3K4me3 peak characteristics of the GS, ZS and ND groups of genes at the four time points, spermatid, sperm, pre-ZGA and ZGA (**Fig.3d**). We observe that GZ genes have significantly higher H3K4me3 peak intensity and peak breadth than all other gene expression groups at all time points, notably also during the transcriptionally quiescent stages (**Fig.3d, Fig.S3e**). The mean peak intensity of GS genes is significantly lower than the GZ group in earlier time points, gradually dropping lower than the mean peak intensity of ZS genes during ZGA **(Fig.3d)**. Groups with zygotically expressed genes i.e., GZ and ZS groups gain asymmetrically broader peaks during ZGA, likely due to active transcription ^11,30^ **(Fig.3d, Fig.S3e)**. This indicates that, in agreement with our hypothesis, genes which are expressed in the gametes and in embryos (GZ group) maintain high intensities and breadth of H3K4me3 marks around their promoters across the interjacent transcriptionally silent phase.

Since the GZ group of genes, which has an active gene expression in both the gametes and the embryo, shows strikingly similar characteristics to the KEPT group of genes, which maintains H3K4me3 from gametes to the embryo, we addressed if they overlap. Indeed, over 80% of GZ genes overlap and are significantly associated with the KEPT group of genes, but not with any of the other groups (**Fig.3e-g; Table S1**). Consistently, the GS group of genes are significantly associated with the LOST group of genes, fitting with their loss of gene expression during development. Genes found to be expressed only in the zygote (ZS group) are associated with the GAINED group of genes, matching their *de novo* activation at ZGA stage. Importantly, however, while almost all GZ genes are also KEPT group of genes, the reverse is not the case – not all KEPT group of genes are also GZ genes **(Fig.3e)**, suggesting that maintenance of H3K4me3 across pre-ZGA stages is not strictly indicative of gene expression at ZGA. Retention of H3K4me3 observed at promoters of some GS genes during ZGA **(Fig.3f)** may possibly allude to zygotic expression of these genes at timepoints outside the window of this study i.e., after 9hpf. On the other hand, H3K4me3 pre-marking at promoters of developmental genes expressed soon after ZGA has been reported zebrafish^7^. Similarly, we wondered whether genes overlapping between ZS and KEPT groups may represent a group of genes required during early development. Together, this suggests that genes that are actively expressed in both the gamete and the embryo also maintain H3K4me3 around their promoters from gametes to early embryos.

Our results hence reveal an association between the genes that maintain H3K4me3 throughout early development and the genes that are transcribed in both early and late stages i.e., spermatid and ZGA, raising the possibility that H3K4me3 is driving the maintenance of active chromatin states.

### Genes maintaining H3K4me3 are expressed early during zygotic genome activation

H3K4me3 around the promoter regions of genes and in the gene body has been reported to be important for high processivity of RNA pol II and consequently with fast and efficient gene transcription^19^. Thus, we hypothesized that pre-marking of genes with H3K4me3 prior to ZGA could allow for particularly early and fast activation of their gene expression during ZGA in *Xenopus* embryos.

We therefore analyzed nascent RNA-seq data^31^ and classified zygotically expressed genes – which include all genes from both the GZ and ZS groups – into four groups based on the timepoint at which their transcripts were first detected **(Fig.3h)**. Indeed, the earliest identified transcripts at 6 hours post-fertilization (hpf) are from genes that are significantly associated with the KEPT group of genes, while the later expressed genes (7hpf, 8hpf, 9hpf) are associated with both KEPT and GAINED genes **(Fig.3i)**. Similarly, the earliest expressed genes (6hpf) are associated with the GZ group of genes, and we observe that the significance of association is progressively less over time **(Fig.3j)**. On the other hand, genes with later detected transcripts (7hpf, 8hpf, 9hpf) are associated with the ZS group of genes, with the significance of association increasing with time **(Fig.3k)**.

When addressing H3K4me3 peak characteristics around the promoters of the ZGA gene groups, we find that genes induced at 6hpf have stronger promoter peak intensities than all other groups at gametes, pre-ZGA and ZGA stages **(Fig.3k)**. Later expressed genes (7-9hpf) gain H3K4me3 asymmetrically from their promoter region towards the 3’ end as ZGA begins **(Fig.3k, Fig.S3f)** and peak breadth increases in parallel, reminiscent of the trend seen for gene promoters of the GAINED group **(Fig.2e, Fig.2f)**. Gene Ontology enrichment analysis showed that the later expressed gene groups enrich for development, regulation and signaling related terms, indicating that these genes are important for supporting the embryo through early development **(Fig.S3g)**.

In summary, we have uncovered that there is a close correlation between H3K4me3 dynamics and gene expression dynamics in the early embryo. Additionally, we find that H3K4me3 is specifically maintained around promoter regions of genes that were transcribed in the gametes, became silent across multiple cell divisions and were then reactivated early during ZGA.

### Maintenance of H3K4me3 during early embryonic cell divisions is required for subsequent proper zygotic genome activation

Having established that H3K4me3 is maintained at the promoter regions of genes through multiple cell divisions when transcription is paused, we next investigated whether this maintenance not only coincides with, but is also functionally linked to, gene expression dynamics at ZGA. Specifically, we aimed to determine whether disrupting the propagation of H3K4 methylation in the cell cycles preceding ZGA, but not during ZGA itself, affects the timely activation of gene expression at ZGA.

Hence, we set up an assay to disrupt H3K4 methylation levels in the embryo within the time window between fertilization and ZGA. We reduced H3K4 methylation orthogonally by overexpressing mRNA encoding the H3K4-specific demethylase Kdm5b^wt^ to remove the methyl residues. We combined this with expression of the H3.3 histone variant with a lysine -to-methionine mutation H3.3^K4M^, which acts in a dominant negative manner to sequester H3K4 methyltransferases and prevent the deposition of new methyl residues^32–34^. The catalytically inactive demethylase Kdm5b^ci^ combined with wild type H3.3^wt^ served as control treatment. All mRNA transcripts were equipped with a C-terminal 3xHA tag and an auxin-inducible degron (AID) sequence to enable their degradation at the 256-cell stage. In this way, interference with H3K4 methylation specifically at pre-ZGA stages was achieved in the treatment condition specifically at pre-ZGA stages.

Thus, a combination of Kdm5b^wt^ and H3.3^K4M^ mRNAs, or a combination of their catalytic inactive counterparts Kdm5b^ci^ and H3.3^wt^ as the control, were co-injected with TIR1 – which degrades AID-fused proteins upon addition of auxin – into embryos at the one-cell stage **(Fig.4a)**. We confirmed expression of the injected mRNAs before ZGA, resulting in 25% depletion of H3K4me3 in pre-ZGA embryos expressing Kdm5b^wt^ and H3.3^K4M^ **(Fig.4b, Fig.4c)** compared to the control conditions. After 8 cell divisions, degradation of overexpressed proteins was induced by addition of auxin, which was found to be complete within 2 hours **(Fig.4b)**. Using immunofluorescence, we confirmed that at mid-ZGA, H3K4me3 levels in these embryos reaccumulate to levels comparable to those in control and wild type embryos **(Fig.4d, Fig.4e)**. In embryos in which Kdm5b^wt^ and H3.3^K4M^ degradation was not induced, H3K4me3 levels remained significantly reduced at mid-ZGA compared to control conditions **(Fig.4d, Fig.4e)**. Thus, our setup successfully reduces H3K4me3 levels within a time window limited to pre-ZGA development.

**Figure 4.**
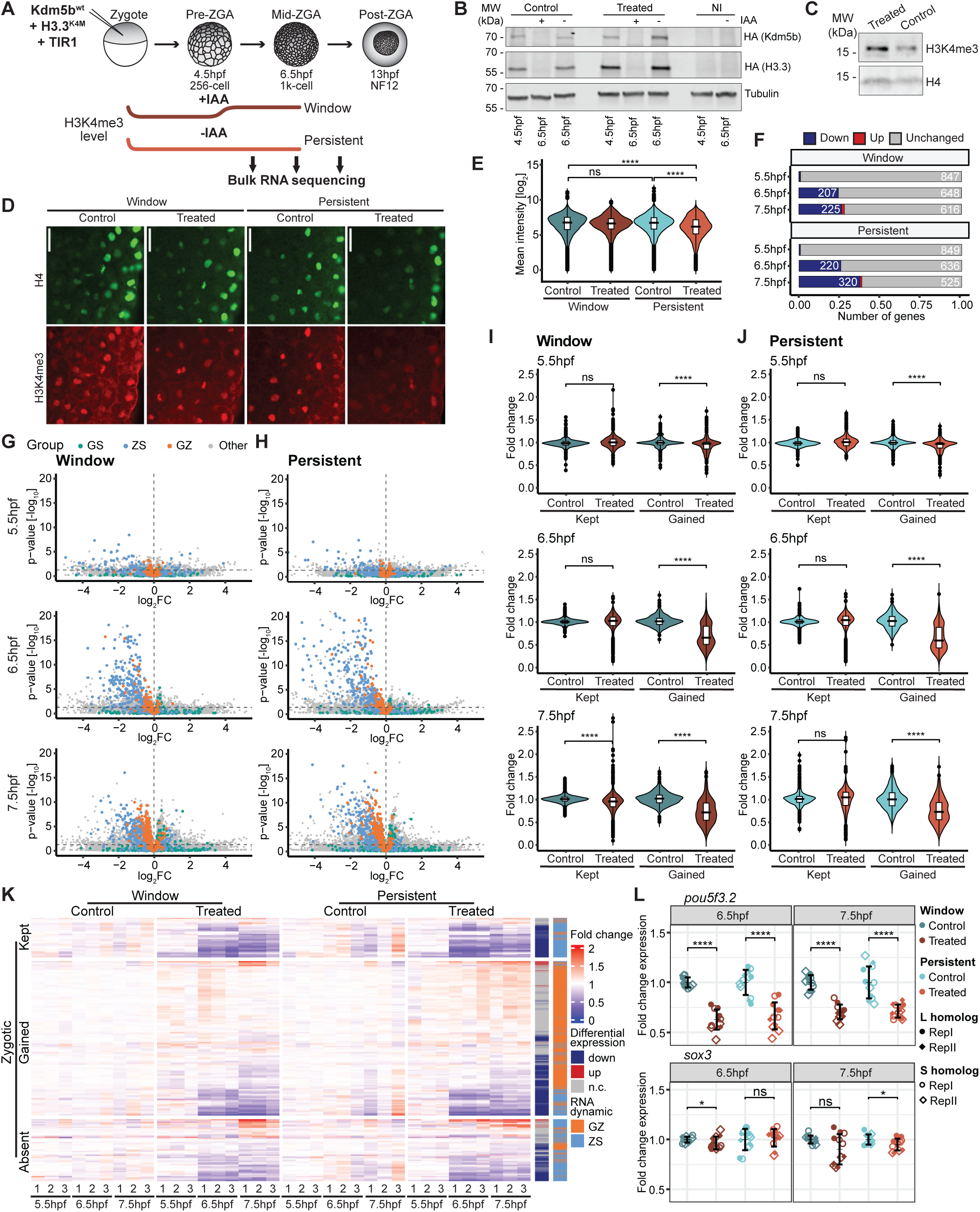
Maintenance of H3K4me3 during early embryonic cell divisions is required for proper ZGA. (A) Illustration of experimental setup for time-specific and persistent H3K4me3 depletion. (B) Western blot showing translation of ectopic HA-tagged Kdm5b and H3.3 mRNA in pre-ZGA embryos and degradation of proteins after IAA induction in cytoplasmic fraction of mid-ZGA embryos, compared to tubulin loading control. Control embryos show Kdm5bci and H3.3wt expression and treated embryos show Kdm5bwt and H3.3K4M expression. “NI” denotes non-injected control embryos. (C) Western blot of pre-ZGA embryos after nuclear isolation showing depletion of H3K4me3 compared to H4 in treated and control embryos. (D) Representative images of H3K4me3 levels in control and treated embryos at mid-ZGA, with or without auxin induction. Nuclei are segmented using H4 (green), and H3K4me3 (red) values are measured. Scale bars: 50µm. (E) Quantification of immunofluorescence based on H3K4me3 signal in segmented nuclei. Statistical test: one-sided Wilcoxon rank-sum test; p-values: (****)<=0.0001, (***)<=0.001, (**) <= 0.01, (*)<=0.05; n.s. are p-values > 0.05. (F) Number of differentially expressed and unchanged zygotic genes at 5.5hpf, 6.5hpf and 7.5hpf time points in window treatment condition compared to control embryos in biological replicate 1 (adj. p-value < 0.05). *Zygotic genes = Gamete+Zygotic (GZ) + Zygotic-Specific (ZS).* (G-H) Volcano plots displaying expression fold-change and adj. p-value (cutoff: 0.05) in treated condition compared to control condition at each timepoint colored by RNA dynamics groups for (G) window depletion and (H) persistent depletion. (I-J) Fold-change of gene expression levels of all zygotic genes in each group calculated over mean TPM of respective control technical replicates (I: Window, J: Persistent) at the three time points in biological replicate 1. (K) Expression of zygotic genes in 3 technical replicates of every sample, clustered by H3K4me3 dynamics group (KEPT, GAINED and ABSENT) for biological replicate 1. Fold-change of every gene is calculated over mean TPM of three technical replicates of the control condition of each respective time point. Genes are annotated by differential gene expression, and RNA dynamics group. (L) Fold change expression of *pou5f3.2* and *sox3* calculated over mean TPM of respective control technical replicates at two timepoints.

Next, we tested whether this temporary disruption of H3K4me3 during early embryonic cell division affects proper onset of ZGA. We treated embryos as described above **(Fig.4a)** but this time analyzed them at three stages spanning the onset of ZGA (5.5hpf, 6.5hpf, 7.5hpf) by RNA-seq. When comparing embryos with perturbed H3K4me3 propagation against control embryos, we detected significant up- and downregulation of transcripts. The number of differentially expressed genes (DEGs) increased with time and significantly overlapped (Fisher’s exact test p-values < 0.0001) across both biological replicates **(Fig.S4a, Fig.S4b)**. A slightly higher number of misregulated genes was identified in embryos where interference with H3K4me3 was persistent **(Fig.S4c)**.

Focusing on all genes found to be expressed at ZGA (see GZ+ZS groups of genes, **Fig.3a)**, we observed that approximately 25% of zygotic genes were significantly downregulated, while almost no zygotically expressed genes were upregulated by our treatments, compared to controls **(Fig.4f, Fig,4g, Fig.S4d, Fig.S4f)**. Importantly, we found that genes which maintain H3K4me3 around their promoters in normal embryos (KEPT group) were strongly affected by these treatments. Mean expression levels of the KEPT group of genes were significantly reduced in treated embryos at 6.5hpf and 7.5hpf compared to controls **(Fig.4i, Fig.S4g)**. In line with this, we saw that downregulated zygotic genes contained many genes that show maintenance of H3K4me3 around their promoters (KEPT group; **Fig.4k, Fig.S4i**), suggesting a direct link between H3K4me3 maintenance prior to ZGA and accurate activation of these genes during ZGA. More unexpectedly, we observed that genes that gain H3K4me3 at their promoters around ZGA stage (GAINED group) were also affected by our treatments: their mean expression levels, too, were lower in the treated embryos compared to controls at 6.5hpf and 7.5hpf **(Fig.4i, Fig.S4g)**. In addition, zygotic genes from the GAINED group were also found among the downregulated genes **(Fig.4k, Fig.S4i)**. Together, this indicated that a temporary disruption of H3K4me3 maintenance affects reliable expression of genes that are accompanied by de novo H3K4me3 deposition at their promoters during ZGA. In addition, we also observed downregulated zygotic genes from the ABSENT group, possibly due to secondary effects of the treatment **(Fig.4k, Fig.S4i)**. Indeed, amongst the downregulated genes belonging to the KEPT group of genes is *pou5f3.2*, corresponding to a transcription factor which, together with Sox3, is known to facilitate ZGA in *Xenopus laevis* **(Fig.4l)**^35,36^. We found that s*ox3* levels are also reduced in embryos subjected to the window treatment when compared to controls, albeit not significantly.

Overall, the persistent reduction of H3K4me3 showed a similar, and only slighter pronounced phenotype than the shorter window treatment preceding ZGA **(Fig.4h, Fig.4j, Fig.S4c, Fig.S4e, Fig.S4h)**. We observed significant downregulation of both transcription factors shown to be important for ZGA, Pou5f3.2 and Sox3, in embryos with persistently reduced H3K4me3 levels **(Fig.4l)**. This suggests that H3K4me3 maintenance prior to ZGA and H3K4me3 acquisition during ZGA-stage are important for accurate activation of gene expression of many zygotic genes.

In summary, our experiments show that restricting H3K4 methylation in the globally transcriptionally silent window before ZGA affects proper expression of both zygotic genes that maintain H3K4me3 during this window and those that gain H3K4me3 during H3K4me3.

### H3K4me3 maintenance around promoters correlates with increased chromatin accessibility, high GC content and DNA hypomethylation

Having established a functional link between proper ZGA and the maintenance of H3K4 methylation from the gametes to the embryo, we next aimed at gaining further insights into the mechanisms underlying H3K4me3 propagation across several transcriptionally silenced cell divisions. In embryonic stem cells (ESCs), connections between GC content, DNA hypomethylation and H3K4me3 deposition mechanisms have been identified^37^. Since we demonstrated that H3K4me3-marked promoters in pre-ZGA embryos are found in distinct chromatin states regarding DNA methylation, GC content and chromatin accessibility **(Fig.1k)**, we explored next whether the patterns of these features vary based on the specific H3K4me3 dynamics and gene expression dynamics that we have classified above.

We focused on the groups of genes that keep, lose or gain H3K4me3 in development. We found that the promoters of the KEPT group of genes have significantly increased DNA accessibility, lower DNA methylation and higher GC content around their promoters than all other groups in gametes **(Fig.S5a)** and in pre-ZGA embryos **(Fig.5a-d)**. Interestingly, genes that gain H3K4me3 at ZGA overall showed lower chromatin accessibility and GC content than KEPT genes, and at the same time higher levels of DNA methylation in pre-ZGA embryos **(Fig.S5c, Fig.S5d)**. These findings indicate that the persistent presence of H3K4me3 around GC rich and accessible promoters during early development is anticorrelated with DNA methylation prior to ZGA.

**Figure 5.**
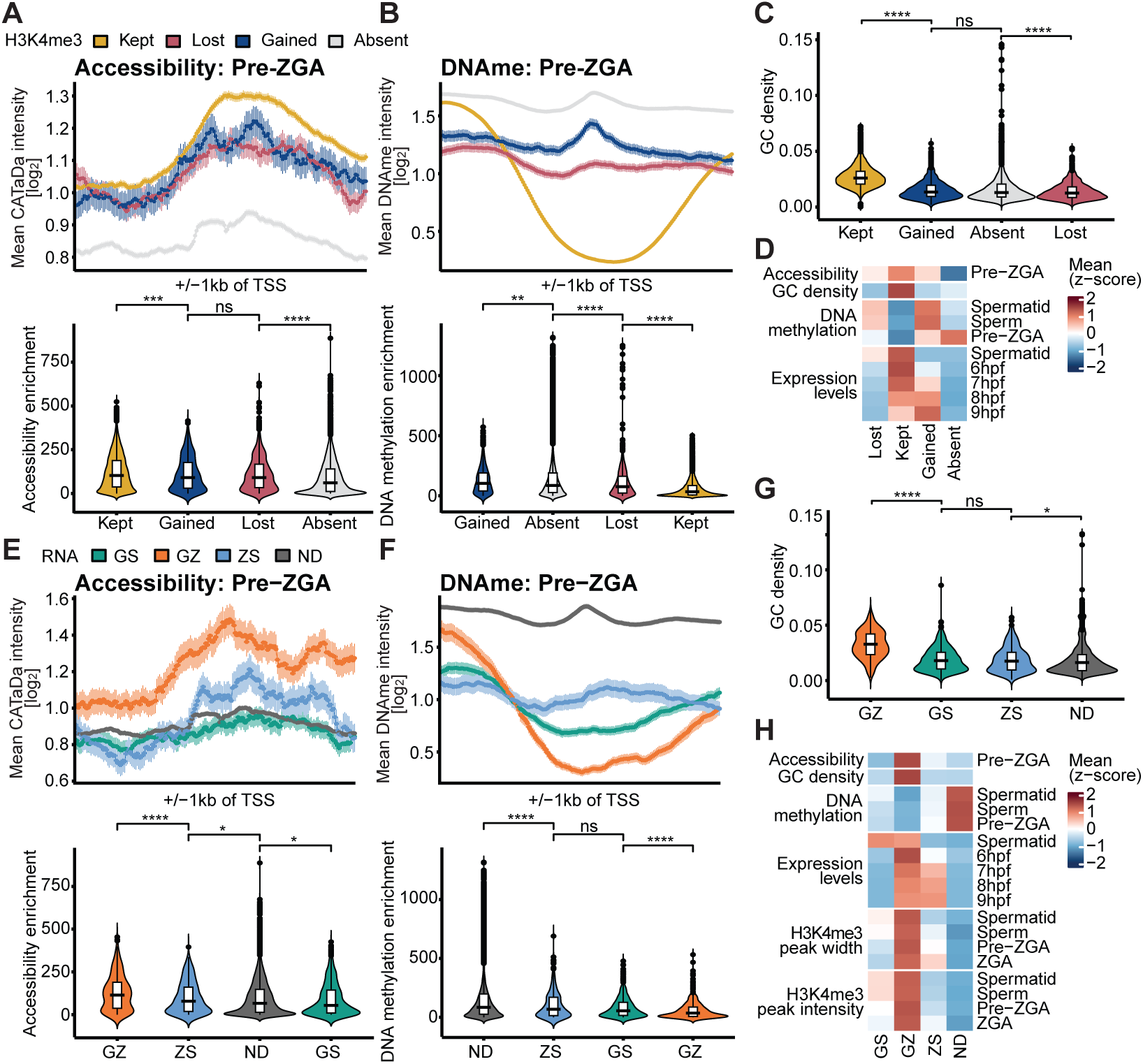
H3K4me3 maintenance around promoters correlates with increased chromatin accessibility, high GC content and DNA hypomethylation. (A) (Top) Accessibility measured by DamID intensity around TSS (+/−1kb) for H3K4me3 dynamics groups at pre-ZGA stage. (Bottom) Accessibility promoter enrichment in each group for pre-ZGA stage. Statistical test: one-sided Wilcoxon rank-sum test; p-values: (****)<=0.0001, (***)<=0.001, (**) <= 0.01, (*)<=0.05; n.s. are p-values > 0.05. (B) (Top) DNA methylation around TSS (+/−1kb) for H3K4me3 dynamics groups for pre-ZGA stage. (Bottom) DNA methylation promoter enrichment in each group for pre-ZGA stage. (C) Comparison of promoter CG density for H3K4me3 dynamics groups. (D) Heatmap showing the z-scored mean of accessibility, CG density and DNA methylation at promoters and expression levels for available timepoints for the major H3K4me3 dynamics groups. (E) (Top) Accessibility measured by DamID intensity around TSS (+/−1kb) for RNA dynamics groups at pre-ZGA stage. (Bottom) Accessibility promoter enrichment in each group for pre-ZGA stage. (F) (Top) DNA methylation around TSS (+/−1kb) for RNA dynamics groups for pre-ZGA stage. (Bottom) DNA methylation promoter enrichment in each group for pre-ZGA stage. (G) Comparison of promoter CG density for RNA dynamics groups. (H) Heatmap showing the z-scored mean of accessibility, CG density and DNA methylation at promoters and expression levels for available timepoints for the different RNA dynamics groups.

Furthermore, the genes in the GZ gene set, which are transcribed in gametes and in embryos and enriched for H3K4me3, also showed high DNA accessibility, high GC content and DNA hypomethylation of their promoters at pre-ZGA **(Fig.5e-h, Fig.S5b)** when compared to other gene expression groups. Promoters of genes whose transcription is first initiated at ZGA (ZS group) showed lower chromatin accessibility, GC content and increased DNA methylation compared to the promoters of the GZ group of genes at the pre-ZGA timepoint **(Fig.5h)**.

Finally, we found that promoters of early expressed genes tend to have higher levels of CpG density **(Fig.S5e, Fig.S5f)**. However, in contrast to H3K4me3 maintenance, chromatin accessibility or DNA methylation, CpG levels of promoters prior to ZGA do not correlate with the timing of gene expression during ZGA **(Fig.S5g-i).**

In summary, we observed that promoters maintaining H3K4me3 and a memory of previously active gene expression states are characterized by high CpG density, low DNA methylation and high accessibility in transcriptionally silent pre-ZGA embryos. This suggests that H3K4me3 maintenance, specifically at promoters of genes that were previously transcribed in the gametes, silenced across multiple cell divisions and then reactivated early during ZGA, is sustained through the help of DNA sequence-specific, DNA hypomethylation-related H3K4me3 reader-writer mechanisms.

### Kmt2b and Cxxc1 facilitate transcription-independent maintenance of H3K4me3 and zygotic gene expression

Previous studies in mouse ESCs and mouse oocytes have identified CFP1 and MLL2 respectively, as H3K4 methyltransferases that can function independently of transcription^38,39^. Both proteins contain a CXXC domain, which reads and recruits the methyltransferase to unmodified CpGs for H3K4 methylation. Interestingly, *Xenopus* CFP1 and MLL2 orthologs *cxxc1* and *kmt2b* were found to be expressed at strikingly high levels during pre-ZGA^28^ **(Fig.S6a, Fig.S6b)**. Thus, we tested if Cxxc1 and Kmt2b contribute to H3K4 methylation in early *Xenopus* embryos and if they are essential for establishing correct gene expression patterns during ZGA.

For this purpose, we injected translation-blocking antisense morpholinos (asMOs) against both homologs of *kmt2b* and *cxxc1*, or control morpholinos (ctrlMO), into one-cell embryos and allowed them to develop **(Fig.6a)**. Although maternally deposited protein is not targeted by this approach, we still observed global depletion of Cxxc1 and Kmt2b protein and, importantly, a corresponding reduction in H3K4me3 levels in asMO-treated embryos when compared to control treated ones **(Fig.6b, Fig.S6c)**.

**Figure 6.**
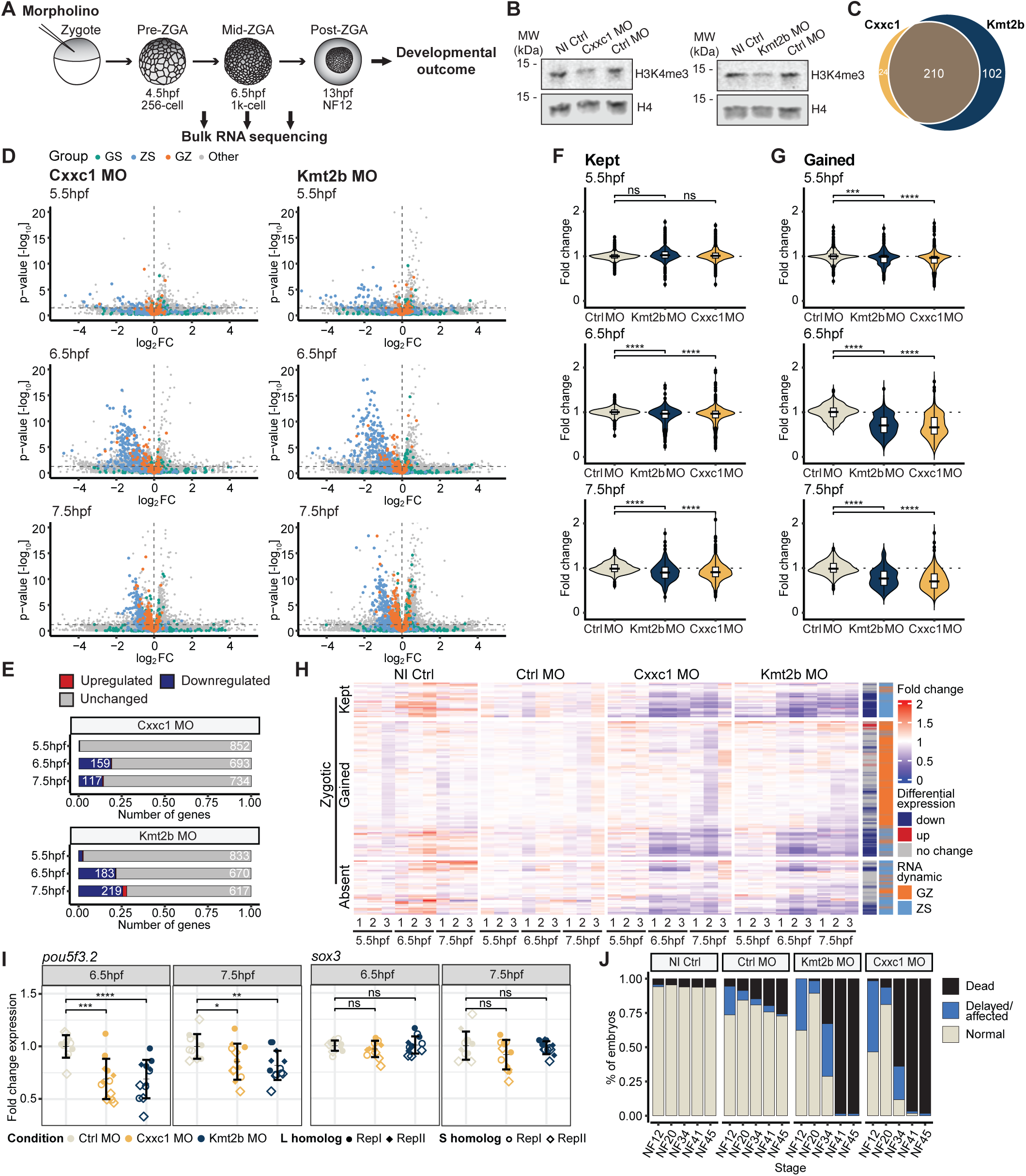
Cxxc1 and Kmt2b are required for proper ZGA and embryonic development. (A) Illustration of experimental setup. (B) Western blot showing H3K4me3 levels in Cxxc1 and Kmt2b knockdown embryos compared to non-injected control and control morpholino-injected embryos at post-ZGA (st12). (C) Overlap of downregulated zygotic genes at any timepoint between Cxxc1 and Kmt2b morpholino conditions compared to control morpholino. (D) Volcano plots displaying expression fold-change and adj. p-value (cutoff: 0.05) in knockdown conditions compared to control morpholino at each timepoint colored by RNA dynamics groups. (E) Number of differentially expressed and unchanged zygotic genes at 5.5hpf, 6.5hpf and 7.5hpf time points in Cxxc1 and Kmt2b morpholino-injected conditions compared to control morpholino embryos (adj. p-value < 0.05). *Zygotic genes = Gamete-Zygotic (GZ) + Zygotic-Specific (ZS).* (F-G) Fold-change of gene expression levels of all zygotic genes in each group calculated over mean TPM of control morpholino technical replicates for (F) KEPT and (G) GAINED groups at the three time points. Statistical test: one-sided Wilcoxon rank-sum test; p-values: (****)<=0.0001, (***)<=0.001, (**) <= 0.01, (*)<=0.05; n.s. are p-values > 0.05. (H) Expression of zygotic genes in 3 technical replicates of every sample, clustered by H3K4me3 dynamics group (KEPT, GAINED and ABSENT). Fold-change of every gene is calculated over the mean TPM of three technical replicates of control morpholino of the respective time point. Genes are annotated by differential gene expression, and RNA dynamics group. (I) Fold change expression of *pou5f3.2* and *sox3* calculated over mean TPM of control morpholino replicates at two timepoints. (J) Quantification of survival phenotype. n=2 experiments, N>35 embryos per condition.

We then investigated the effect of Cxxc1 and Kmt2b knockdown, and concomitant H3K4me3 depletion, on ZGA. We performed RNA sequencing on Cxxc1 and Kmt2b knockdown embryos at three stages spanning the onset of ZGA (5.5hpf, 6.5hpf, 7.5hpf; **Fig.6a**). We then performed differential gene expression analyses and observed significant up- and downregulation of transcripts in both Cxxc1 and Kmt2b knockdown embryos, when compared to the controls, with the number of differentially expressed genes (DEGs) increasing with developmental time **(Fig.S6d)**. We identified significant overlap (Fisher’s exact test p-value < 0.0001) of downregulated genes across Kmt2b and Cxxc1 knockdown conditions **(Fig.6c, Fig.S6e)** and biological replicates **(Fig.S6f)**. We found that downregulated genes in both conditions contain zygotically expressed genes of the GZ and ZS groups **(Fig.6d, Fig.6e, Fig.S6g)**. In contrast, upregulated genes consist mainly of non-zygotic genes **(Fig.6d)**. This indicates deregulation of accurate ZGA.

We therefore focused our analyses on zygotic genes, classified earlier in our analyses (GZ and ZS groups of genes, see **Fig.3a**), and found that approximately 20% of these were downregulated in Cxxc1 knockdown embryos, and roughly 25% were downregulated in Kmt2b knockdown embryos by 7.5hpf **(Fig.6e, Fig.S6g)**. Importantly, out of these downregulated zygotic genes, genes belonging to the groups that keep H3K4me3 around their promoters during early embryonic development (KEPT group) and genes that gain H3K4me3 during ZGA stage (GAINED group) were affected in Cxxc1 and Kmt2b knockdown embryos. With increasing developmental time at 6.5hpf and 7.5hpf timepoints, we observed significant downregulation of genes in the KEPT and GAINED groups in Cxxc1 or Kmt2b knockdown versus control embryos **(Fig.6f-h, Fig.S6h-j)**. Additionally, we observed a significant overlap of downregulated zygotic genes in individual Cxxc1 and Kmt2b knockdown conditions, and both conditions affect expression of developmental genes and genes involved in transcription regulation and developmental processes **(Fig.S6k)**. *Pou5f3.2* and s*ox3* were amongst the downregulated genes in the KEPT-group: In both Cxxc1 and Kmt2b asMO-injected embryos, we observed significant reduction of *pou5f3.2* expression levels when compared to controls, but only a slight, non-significant reduction in s*ox3* levels (**Fig.6i**). Together, these observations link Cxxc1 and Kmt2b to H3K4me3 propagation and acquisition during embryonic development in *Xenopus* and demonstrate a role for these enzymes in proper ZGA in *Xenopus*.

We then addressed whether the observed depletion of Kmt2b and Cxxc1 in embryos was affecting embryonic development. Both Cxxc1 and Kmt2b knockdown embryos showed a delay in development compared to control morpholino and wildtype embryos. Similar defects, including aberrant gut formation and an overall twisted and shorter body axis, as well as a “bent tail” phenotype affecting swimming ability, could be observed in tadpoles of both knockdown conditions but not in control embryos **(Fig.S6l)**. Tadpoles with methyltransferase knockdown did not survive past the feeding tadpole stage (4 days post fertilization), demonstrating that Cxxc1 and Kmt2b functions are each individually essential for proper embryonic development **(Fig.6j)**.

Thus, our results reveal that the methyltransferases Kmt2b and Cxxc1 regulate accurate activation of zygotic genes that either maintain or gain H3K4me3 during embryonic development. Furthermore, we show that these H3K4me3 reader-writer enzymes acting independently of transcription are important for faithful establishment of zygotic gene expression patterns and embryonic development.

## Discussion

Our study identifies H3K4 methylation-mediated memory of active chromatin states and reveals its crucial role in proper zygotic genome activation and embryonic development. We provide evidence that H3K4me3 is established around gene promoters in gametes and maintained in embryos across multiple cell divisions despite global repression of transcription. We show that proper maintenance of H3K4me3 during these stages is crucial to ensure the establishment of correct embryonic gene expression patterns during ZGA. H3K4me3 propagation is facilitated by the activity of H3K4 methyltransferases Cxxc1 and Kmt2b at hypomethylated, CpG-dense gene promoters. Interference with these enzymes results in reduced H3K4me3 levels, aberrant zygotic genome activation and developmental failure. Therefore, our study demonstrates that H3K4me3 maintenance is critical for proper ZGA and early embryonic development, with transcription-independent H3K4 methyltransferases playing a key role in this process.

By subjecting histones of pre-ZGA embryos to mass spectrometry analyses, we uncovered the presence of numerous histone modifications on pre-ZGA chromatin, including active histone modifications. This is earlier than previously anticipated and is unexpected, given the highly dynamic nature of early embryonic chromatin: It is replicated on average every 30 minutes in the absence of transcription. We find that H3K4me3 is nevertheless maintained with the help of the transcription-independent methyltransferases Kmt2b and Cxxc1 around promoter regions of a large proportion of genes. Our findings reveal a novel role for Kmt2b in maintaining canonical promoter H3K4 methylation during the transcriptionally silent and rapidly dividing pre-ZGA phase in *Xenopus laevis* embryos, contrasting with its reported role in acquiring non-canonical broad H3K4me3 domains in non-diving transcriptionally active mouse oocytes^39^. We envision that these enzymes recognize the GC-rich, hypomethylated promoter regions of these genes and reinstate H3K4me3 after DNA replication. Broad H3K4me3 domains spanning several nucleosomes correlate with effective maintenance of the mark, possibly by increasing the chance of reestablishing of the chromatin state in the short time window after replication and before entering the next cell cycle. It is possible that, in addition, not yet determined transcription factors and DNA sequence elements contribute to H3K4me3 maintenance. Importantly, however, our data suggests that H3K4me3 maintenance occurs independently of transcription and of the signal that initially induced the state during gamete maturation. While interference methods currently cannot distinguish between the contribution of H3K4 di- and trimethylation, we can conclude that loss of H3K4 methylation results in defective gene activation and embryonic death. Thus, together, these results point towards H3K4 methylation being a mechanism of epigenetic memory of active chromatin states, supporting our previous findings in nuclear transfer experiments^13^. It will be interesting to determine if other active histone modifications detected on pre-ZGA chromatin such as H3K9ac, H3K27ac and H3K4me1 (**Fig.1b**) also contribute to epigenetic memory and successful embryonic development.

H3K4me3 and DNA methylation have long been studied as mutually antagonistic factors, as H3K4me3 impedes DNA methylation by inhibiting DNMT3 binding, responsible for *de novo* DNA methylation in mouse^24,40^. Conversely, hypomethylated DNA can recruit certain H3K4 methyltransferases via the CXXC domain^38,39,41^. This raises the question of whether H3K4me3 is maintained in early development as a response to DNA methylation patterns, or whether the observed hypomethylation is a byproduct of consistent H3K4me3 peaks at these promoters. Previous work in *Xenopus tropicalis* has hypothesized that H3K4me3 is absent in cleavage-stage embryos and is acquired during ZGA primarily based on DNA methylation logic to hypomethylated CpG-dense regions^11^. Our findings challenge and expand upon these conclusions by directly inspecting the chromatin landscape at the pre-ZGA stage. We demonstrate that H3K4me3 is not only present at this stage, but also that it is maintained from gametes to embryos across a transcriptionally silent window. Although our observations of the KEPT and GZ groups of genes show that H3K4me3 maintenance is biased towards regions with hypomethylated, CpG-dense promoters, our interference experiments highlight that that this trend is not merely a byproduct of DNA methylation status. Rather, we show that H3K4me3 maintenance is steered towards gene promoters relevant for successful early embryonic development.

What is the role of H3K4me3 maintenance during early development? In previous experiments in zebrafish and mouse addressing the roles of histone modifications and variants such as H3K4me3, H3K27ac, H3K9me3, H3K27me3, H2AK119ub and H2A.Z in supporting ZGA, embryos were usually subjected to interference from fertilization onwards across ZGA and development^32,42–55^. Unfortunately, this does not allow us to distinguish whether the propagation of the histone mark before ZGA or the accumulation of the mark throughout early development is important for establishing accurate gene expression patterns at ZGA. While we did not specifically address the latter with our experiments, we find that interfering with the propagation of H3K4 methylation in a time window prior to ZGA has a gene expression phenotype almost as strong as persistent depletion throughout early development, including ZGA. This suggests that maintenance of H3K4me3 during early developmental cell divisions with repressed transcription prior to ZGA is important for proper onset of ZGA and successful *Xenopus* embryonic development.

Surprisingly, the canonical H3K4me3 patterns that we have investigated in pre-ZGA *Xenopus* embryos align with the occurrence of mostly canonical H3K4me3 found in human oocytes and embryos^56^, unlike in mouse oocytes^46^. Additionally, Mll2 is expressed in early human embryos and is depleted as ZGA begins, much like the expression patterns of its homolog Kmt2b in *Xenopus* embryos^56^. Taken in context with other species similarities such as the relatively higher degree of paternal histone retention after fertilization in both species and multiple transcriptionally silent cell divisions before ZGA^12,57–59^, *Xenopus* may provide unique insights into the mechanisms supporting epigenetic memory of active states in pre-ZGA human embryos.

On a molecular level, we find that early transcriptional reactivation of genes during ZGA correlates with H3K4me3 maintenance and high GC content, but not with DNA accessibility or DNA methylation. It has been shown previously that H3K4me3 around the promoter region and in the gene body is important for high processivity of RNA Pol II to ensure fast and efficient transcription^19^. Thus, it is tempting to speculate that H3K4me3 maintenance around promoter regions from the gametes and pre-ZGA embryos drives fast and efficient expression of these genes early during ZGA. The products of these genes, which are enriched for housekeeping functions, likely promote further effective ZGA. Interestingly, amongst those genes are also the two transcription factors reported to facilitate ZGA in *Xenopus*, the maternal pluripotency factors Sox3 and Pou5f3.2^35,36^. Combinatorial knockdown of these factors in *Xenopus* causes developmental arrest at gastrulation^35,36^. Both the persistent and the brief window disruption of H3K4me3 maintenance prior to ZGA result in a significant downregulation of transcripts encoding Pou5f3.2 and, to a lesser extent, Sox3. A consequence of this may be decreased protein levels of Sox3 and Pou5f3.2, reducing the efficiency of their pioneer activity and causing further deregulation of ZGA^35^.

In summary, we propose that H3K4me3 drives the memory of active chromatin states from gametes to embryos by maintaining its presence around promoter regions of genes across multiple cell divisions, even when transcription is repressed. Full H3K4 methylation maintenance is ensured by the methyltransferases Cxxc1 and Kmt2b at hypomethylated, CpG-dense promoters of genes. This maintenance is crucial for ensuring proper zygotic gene activation (ZGA) and successful embryonic development.

## Materials and methods

### *Xenopus laevis* culture

Adult *Xenopus Laevis* were obtained from Nasco (901 Janesville Avenue, P.O. Box 901, Fort Atkinson, WI 53538-0901, USA) and Xenopus1 (Xenopus1, Corp. 5654 Merkel Rd. Dexter, MI. 48130). All frog maintenance and care were conducted according to the German Animal Welfare Act. Research animals were used following guidelines approved and licensed by ROB-55.2-2532.Vet_02-23-126.

### Embryo collection

Sexually mature females were pre-primed for ovulation by injection of 150 IU of Ovogest 300 I.E./ml into their dorsal lymph sac 4-5 days prior to experiments. 500 IU of the same hormone at 1000 I.E./ml was injected on the day before experiments and the females were kept overnight at 14°C in 1x MMR (Marc’s Modified Ringers buffer: 100mM NaCl, 2mM KCl, 1mM MgSO4, 2mM CaCl2, 0.1mM EDTA, 5mM HEPES pH 7.8). Females were then moved to room temperature to lay eggs. Eggs were collected timely and *in vitro* fertilization was performed.

### *In vitro* fertilization

*In vitro* fertilization was performed on eggs by shredding a piece of freshly dissected testes between the two arms of a riffled pair of forceps to release the sperm in a glass dish containing eggs in a minimal amount of 1x MMR. After a 2-minute incubation, the 1x MMR was diluted approximately 10-fold with distilled water and the fertilized eggs were kept at room temperature for 20-30 minutes. Subsequently, the jelly coating on the fertilized eggs was chemically removed using a dejellying solution (2% L Cysteine in milliQ H2O, pH 7.8; adjusted by NaOH), followed by washes in 0.1x MMR.

### Microinjections

For labeling nascent RNA, embryos were injected with 9.2nl of 50µM 5-EU at the 1-cell stage using a NANOINJECT II microinjector (Drummond Scientific Company). For transcription inhibition, embryos were co-injected with 4.6ng ⍺-amanitin. For all injection experiments, embryos were kept in 0.5x MMR during and up to 2 hours post-injections, after which they were transferred to 0.1x MMR for further development.

### Nuclear isolation of pre-ZGA embryos

200 256-cell stage embryos per sample were collected at the desired stage. Buffer was removed and embryos were gently washed thrice using embryo extraction buffer (10mM HEPES pH 7.7, 100mM KCl, 50mM Sucrose, 1mM MgCl2, 0.1mM CaCl2) and then spun down for 1 minute at 700g. All excess liquid was removed, and samples were stored at −80°C until further processing. Frozen embryos were thawed on ice and spun at 17,000 g for 10 min at 4°C. This results in the formation of 3 layers, the upper lipid layer and bottom layer of cellular debris are avoided, and the middle grey layer of nuclei is collected using a P200 pipette. The nuclear fraction is washed twice with SuNASP buffer (250 mM Sucrose, 75mM NaCl, 0.5mM Spermidine, 0.15mM Spermine) with a 5-minute spin at 4°C at 3500g and then twice with RIPA buffer (50mM Tris-HCl pH 7.4, 1% NP40, 0.25% Na-deoxycholate, 150mM NaCl, 1mM EDTA, 0.1% SDS, 0.5mM DTT, 5mM Na-butyrate) supplemented with protease inhibitors, spun for 10 minutes at 14,000 g at 4°C. A Kimwipe is used to aid the removal of yolk and debris caught on the walls of the tube while removing the supernatant between washes. For SDS-PAGE, the pellet was resuspended in 2x E1 buffer (50mM HEPES-KOH pH 7.5, 140mM NaCl 0.1mM EDTA pH 8.0, 10% Glycerol, 0.5% Igepal CA-630, 0.25% Triton X-100, 1% β-mercaptoethanol) and 4x Laemmli sample buffer (BIO-RAD, #1610747) supplemented with 10% β-mercaptoethanol. An equivalent of 50 embryos was loaded per well.

### Western Blotting

For testing histone modifications in gastrula (NF stage 12) embryos, 1-3 embryos were collected and resuspended in E1 buffer followed by centrifugation at 4°C for 2 minutes at 3500 rpm. The supernatant containing the cytoplasmic fraction was collected in a separate tube and the pellet was dislodged with a pipette tip, resuspended in Tris/SDS solution (50mM Tris-HCl pH6.8 + 2% SDS) supplemented by protease inhibitors and vortexed well. 4x Laemmli buffer was added to both chromatin and non-chromatin bound fractions, chromatin was fragmented by manual disruption and heated for 15 minutes at 95°C. 1/4th of an embryo was loaded per well.

For testing cytoplasmic proteins in early cleavage stage embryos, embryos were resuspended in RIPA buffer (50mM Tris-HCl pH 7.4, 1% NP40, 0.25% Na-deoxycholate, 150mM NaCl, 1mM EDTA, 0.1% SDS, 0.5mM DTT, 5mM Na-butyrate) supplemented with protease inhibitors and spun down at 14,000 g for 10 minutes at 4°C. The supernatant containing the cytosolic fraction was collected and 4x Laemmli buffer was added, samples were manually disrupted and vortexed well, followed by heating at 95°C for 15 minutes. 1/4th of an embryo was loaded per well.

SDS-PAGE was performed on 4-20% gradient gels from (Bio-Rad, Mini-PROTEAN TGX #4561096) followed by transfer on PVDF membranes and blocking in 5% BSA in TBS. Depending on the experiment, antibodies were used at 1:1000 dilutions and were incubated overnight at 4°C (see table 3 for antibody details). Membranes were imaged using the Licor Odyssey CLx system and image analysis was performed using ImageJ.

### Histone tail mass spectrometry and data analysis

Pre-ZGA (256-cell stage i.e. 4.5 hpf) and mid-ZGA (4000-cell stage i.e. 6.5 hpf) embryos were prepared using the nuclear isolation method as described above. An equivalent of 50x embryos was loaded per well on a 4-20% gradient gel for SDS-PAGE to resolve histones. Gels were stained using a Coomassie R-250 staining solution (0.2% Coomassie R-250 (w/v), 30% ethanol, 10% glacial acetic acid) at room temperature for 30-40 minutes. Gels were then destained (30% ethanol, 10% glacial acetic acid) at room temperature every few hours for approximately 24 hours until distinct histone bands are achieved against a transparent background. Gels were excised between the 10kDa-20kDa range and in-gel sample preparation was performed for mass spectrometry.

Gel pieces containing histones were washed with 100mM ammonium bicarbonate, dehydrated with acetonitrile, chemically propionylated with propionic anhydride and digested overnight with trypsin. Tryptic peptides were extracted sequentially with 70% acetonitrile/0.25% TFA and acetonitrile, filtered using C8-StageTips, vacuum concentrated and reconstituted in 15µl of 0.1% FA.

For LC-MS/MS purposes, desalted peptides were injected in an Ultimate 3000 RSLCnano system (Thermo) and separated in a 25-cm analytical column (75µm ID, 1.6µm C18, IonOpticks) with a 50-min gradient from 2 to 37% acetonitrile in 0.1% formic acid. The effluent from the HPLC was directly electrosprayed into an Orbitrap Exploris 480 HF (Thermo) operated in data dependent mode to automatically switch between full scan MS and MS/MS acquisition with the following parameters: survey full scan MS spectra (from m/z 250–1200) were acquired with resolution R=60,000 at m/z 400 (AGC target of 3×10^6^). The 15 most intense peptide ions with charge states between 2 and 6 were sequentially isolated to a target value of 2×10^5^ (R=15,000) and fragmented at 27% normalized collision energy. Typical mass spectrometric conditions were: spray voltage, 1.5 kV; no sheath and auxiliary gas flow; heated capillary temperature, 250°C; ion selection threshold, 33.000 counts.

Data analysis was performed with the Skyline (version 21.2) by using doubly and triply charged peptide masses for extracted ion chromatograms. Automatic selection of peaks was manually curated based on the relative retention times and fragmentation spectra with results from Proteome Discoverer 1.4. Integrated peak values were exported for further calculations. The relative abundance of an observed modified peptide was calculated as percentage of the overall peptide.

### Immunofluorescence and Image Analysis

Immunofluorescence and confocal imaging of nascent transcripts in whole-mount embryos was performed as previously described^31,60^ with minor modifications. Briefly, embryos were developed until the desired mid-ZGA stage (i.e. 6.5hpf/1000-cell stage) and fixed using 4% paraformaldehyde (PFA) in 1x MEM (100mM MOPS pH 7.4, 2mM EGTA, 1mM MgSO4) overnight at 4°C, permeabilized in 100% methanol and stored at −20°C until further processing. Embryos were bleached in a solution of 4% formamide and 2% hydrogen peroxide in a 0.5× saline sodium citrate (SSC; 75 mM NaCl, 7.5 mM sodium citrate) buffer for approximately 4-6 hours, until evenly white. 5-EU-injected embryos were incubated with 100mM Tris-HCl, 1mM CuSO4, Alexa Fluor 594 Azide (Thermo Fisher Scientific, #A10270) and 100mM ascorbic acid for 6 hours at room temperature in the dark to stain nascent RNA. Embryos were blocked in 1% BSA in TBST (1x TBS containing 0.1% Triton-X-100, pH 7.6) for 3 hours at RT, followed by primary antibody staining depending at 4°C overnight, with the following antibodies depending on the experiment: ⍺-H4 (1:500; Abcam, ab81380), ⍺-H3K4me3 (1:300; Abcam, ab8580). Secondary antibody incubation and DNA staining was simultaneously performed overnight at 4°C in the dark, with the following antibody fluorescent conjugates or stains depending on the experiment: goat ⍺-mouse AF488 (1:500; Invitrogen, #A-11001), goat ⍺-rabbit AF647 (1:500; Invitrogen, #A-21244), SiR-DNA (1:5000; Tebubio, #SC007). After staining, embryos were post-fixed in 4% PFA in 1x MEM for 2 hours at room temperature, dehydrated in methanol, and cleared in freshly made benzyl alcohol:benzyl benzoate (1:2) solution for 24–48 hours before imaging. All intermediate washes were performed as described earlier^31,60^. Cleared embryos were mounted in chambers constructed from cover slips and double-sided tape, filled with clearing reagent. Imaging was performed using a Nikon Ti-2 microscope equipped with Dragonfly confocal unit with a 20× Plan Apo NA 0.75 objective (Nikon) and a Zyla sCMOS camera (Andor), capturing z-stacks with 5 μm intervals and tiling with 2% overlap. Embryos were imaged from both sides to ensure complete signal capture.

For nuclear segmentation, a custom Cellpose model was trained by random extraction of optical sections from z-stacks using a Fiji macro (ImageJ version 1.54f) followed by manual annotation. Images were then split into training and test datasets and the Cellpose model “cyto3” was used for re-training with a mean diameter of 25 pixels determined from the average size of training masks. The training process lasted 300 epochs (Cellpose version 3.0.8, Python version 3.10.0, Python libraries Pytorch 2.3.1, CUDA 11.8, and cuDNN 8.7). The custom-trained model was used for nuclear segmentation on the respective DNA-staining channel, depending on the experiment. Over-segmented labels were corrected by enlarging objects in the images using the morphological operation “dilation”. Identified masks were applied to other channels to measure intensity within the mask area and the “regionprops” module from the scikit-image library (version 0.23.2) was used to extract a set of parameters as the intensity per volume from the nuclei masks and the raw intensity images. Statistical analysis as well as plotting was performed in R (Version 4.3.1) with the packages ggplot2 (version 3.3.6), ggpubr (Version 0.6.0) and multcomp (Version 1.4-26). Figures for microscopy images were made in ImageJ (Version 1.54f). The analysis pipeline was run on an Intel i9 12900KF 24-Core Processor with 64 GB of RAM and an Nvidia RTX A4500 GPU (driver version 525.78.01).

### MBD-seq sample collection, library preparation and sequencing

250x pre-ZGA (256-cell stage i.e. 4.5 hpf) embryos were collected per replicate and prepared using the nuclear isolation method as described above. Nuclear pellet was resuspended in 200µl 1x TE buffer (10 mM Tris-HCl pH 8, 0.1 mM EDTA) and incubated with 4µl RNase A (Sigma Aldrich, #10109142001) for 1 hour at 37°C. Samples were incubated for 1 hour with 10mg/ml proteinase K (Sigma Aldrich, #P6556) at 65°C, followed by heat-induced proteinase K inactivation by a 15 minute incubation at 95°C. 400µl of phenol/chloroform/isoamyl alcohol (25:24:1, pH 8) was added to the sample and vortexed for 30 seconds, followed by a 3 minute centrifugation at maximum speed for 3 minutes and upper, aqueous layer was collected. This step was repeated once. 100µl chloroform was added, sample was vortexed and centrifuged with the same conditions. Upper phase was transferred to a new tube, taking care to avoid chloroform contamination. Ethanol precipitation was performed by adding 12 μL glycogen (5 mg/mL), 40 μL sodium acetate (3 M, pH 5.2), and 1200 μL ethanol (96%), followed by overnight incubation at −20°C. The sample was centrifuged at 4°C for 10 minutes at maximum speed, followed by two 70% ethanol washes. Pellet was air-dried and eluted in 30µl nuclease-free water. Extracted DNA samples were sonicated with Bioruptor Pico (Diagenode) in 0.1ml microtubes (#C30010015) for 8 cycles (ON: 15s/OFF: 90s). Sonication to ∼400bp fragment length was confirmed on a 0.8% agarose gel. Sample concentration was measured using Qubit and diluted using 1x TE buffer to a final concentration of 100 ng/nl. 1.2µg DNA was used for further processing, of which 10% was kept aside as input. Methylated DNA was captured and eluted from remaining samples in one single elution with 150µl High Elution Buffer using the Methylated DNA Capture kit (Diagenode: #C02020010) as described in their protocol. Samples were then purified using phenol:chloroform:isoamyl alcohol isolation and precipitated overnight at −20°C as described above. Samples were finally eluted in 50µl nuclease-free water and per sample, 10-20 ng of total DNA was processed for library preparation.

Sequencing libraries were generated using the NEBNext Ultra II DNA Library Prep Kit for Illumina (NEB, #E7645S) as per the manufacturer’s instructions using 6-8 PCR amplification cycles. The quality of cDNA libraries was assessed using the Agilent High Sensitivity D5000 ScreenTape System (Agilent, #5067-5592) in the Agilent 4150 TapeStation System. Libraries were multiplexed and sequenced was performed as paired-end 50 bp reads on the Illumina NextSeq 1000 by the Helmholtz Core Facility Genomics (CF-GEN).

### MBD-seq and ChIP-seq data processing

Sequencing reads were aligned to the Xenopus laevis genome (v10.1) with bowtie (v4.2). The alignment process excluded discordant and mixed reads, with paired-end reads mapped to the reference genome. Aligned reads were converted to BAM format and filtered to retain only properly paired alignments. HOMER (v4.11) was used for peak identification against corresponding input controls. Broad peaks were called using histone-style parameters, while narrow peaks were called with default settings. Identified peaks were converted to BED format for downstream analysis.

### CATaDa sample collection, library preparation and sequencing

Chromatin accessibility of pre-ZGA embryos was profiled using CATaDa (Chromatin Accessibility profiling using Targeted DamID) as described previously^26^. Embryos were injected at the 1-cell stage with 2.3ng pRN3P_AID-Dam-only construct (a kind gift by Maria-Elena Torres-Padilla; Addgene plasmid # 136065)^61^. 500x injected or non-injected control pre-ZGA (256-cell stage i.e. 4.5 hpf) embryos were collected per replicate and prepared using the nuclear isolation method as described above. Nuclear pellets were resuspended in 400µl gDNA Tissue Lysis Buffer from the Monarch Genomic DNA Purification Kit (#T3010S) and the instructions for lysis of animal tissue and subsequent genomic DNA binding steps were followed as described in the protocol. Elution was performed with 100µl preheated gDNA Elution Buffer as described. Samples volumes were then decreased to 15µl using a SpeedVac instrument for 25 minutes.

All samples were processed for CATaDa as described previously^62^. CATaDa fragments were prepared for Illumina sequencing according to a modified TruSeq protocol^62^. Sequencing was performed as paired end 50 bp reads by the CRUK Genomics Core Sequencing facility on a NovaSeq 6000.

### CATaDa data processing

Analysis of fastq-files from CATaDa experiments was performed with the damidseq pipeline script^63^. Preprocessed fastq-files were mapped to the Xl.v10.1 genome assembly using bowtie2, and mapped reads were binned into fragments delineated by 5’-GATC-3’ motifs. Files were converted to the bigwig file format with bedGrapgToBigWig (v4) for visualization with the Integrative Genomics Viewer (IGV) (v2.13.0). MACS (v3.0.1)^64^ was used to call broad peaks on the mapped read (.bam) files generated by the damidseq_pipeline.

### Signal intensity calculation for ChIP-seq, MBD-seq and CATaDa

The resulting coverages from the preprocessings explained above were log2 transformed after adding a pseudo count of 1. Final signal intensities for Meta plots and Heatmaps were calculated by adding up the coverage of all replicates.

Meta plots (line plots) – Aggregate line plots, depicting the mean signal intensity of gene sets of interest with bands representing standard error of the mean, were generated using ggplot2 (v3.4.4).

Heatmaps – Heatmaps were generated using the EnrichedHeatmap package (v1.32.0) in R. Each row represents the signal intensity of one gene.

Violin plots – Violin plots show the enrichment calculated with the enriched_score function of the EnrichedHeatmap package within the promoter regions of the genes. Promoter regions were defined as ±1kb windows centered on the TSS. A one-sided Wilcoxon rank-sum test was used to calculate statistical significance. Four stars (****) denote a p-value smaller or equal than 0.0001. Three stars (***) denote a p-value smaller or equal to 0.001. Two stars (**) denote a p-value smaller or equal to 0.01. One star (*) denotes a p-value smaller or equal to 0.05. P-values larger than 0.05 are denoted with n.s..

### H3K4me3 dynamics group definitions

Broad peaks were called on publicly available spermatid, sperm, pre-ZGA and ZGA stage H3K4me3 ChIP-seq datasets^12,28,29^ as described in the ChIP-seq processing section above. H3K4me3 dynamics groups were defined based on the detection of a H3K4me3 peak within the gene promoter regions (TSS +/− 1kb) at each time point. Genes for which a H3K4me3 promoter peak is observed at all four time points are denoted as the “KEPT” group of genes. Genes for which a H3K4me3 peak is not detected in the spermatid stage, but is acquired during one of the subsequent stages are labelled as the “GAINED” group of genes. Similarly, genes for which the promoter H3K4me3 peak is detected at the spermatid stage and no longer detected from one of the subsequent stages are called the “LOST” group of genes. Genes for which we did not detect a H3K4me3 peak at any of the four time points were labelled as the “ABSENT” group of genes. These major H3K4me3 dynamics groups can be further categorized based on the timing at which the H3K4me3 peak is lost or gained i.e., the “LOST” group is subdivided into “Lost at sperm”, “Lost at pre-ZGA” and “Lost at ZGA” and the “GAINED” group is subdivided into “Gained at pre-ZGA” and “Gained at ZGA”. Genes that oscillate between loss and gain of H3K4me3 peaks at their promoters during the four time points were removed from further analysis. Additionally, we removed all genes that gained a new peak at the promoter between the spermatid and sperm stages. Collection of spermatid and sperm chromatin samples involves their careful separation from neighbouring blood vessels using testes tissue homogenization and centrifugation, risking blood cell contamination. Indeed, gene ontology analysis of genes that gain new H3K4me3 promoter peaks in the sperm stage showed enrichment for blood and circulation related terms, suggesting possible contamination of blood DNA in the samples^12^.

### Gene expression group definitions

RNA dynamics groups were defined with the help of published total RNA-seq datasets from spermatid^29^ and egg^28^ stages as well as published nascent RNA-seq datasets for embryos at 5hpf, 6hpf, 7hpf, 8hpf and 9hpf^31^. Alignment of sequencing reads and quantification of transcript abundance was performed as described in the “RNA-seq data processing” section below. First, nascent transcription at each time point (6/7/8/9hpf) was calculated based on the net increase of reads from the “background” pulldown at 5hpf, as described in the reference study^31^. Then, transcripts that could be detected above the defined threshold (TPM increase > 5) in all replicates of any embryonic time point (6/7/8/9hpf) were identified as zygotically expressed genes. This stringent list was split into two groups: (1) the genes that were also detected at the spermatid or egg stages were defined as the “Gamete+Zygotic” group (2) the remaining genes were defined as the “Zygotic-Specific” group. In order to define a strict “Gamete-Specific” group, we identified transcripts that satisfied the following conditions: (1) detected above the threshold (> 5 TPM) in the spermatid and/or egg stages, (2) not detected above the threshold (<= 0 TPM increase) in any replicate of the embryonic time points (6/7/8/9hpf). The “Not Detected” group is defined by all genes that are not detected above the threshold (<= 0 TPM increase) in any stage (spermatid, egg, 6/7/8/9hpf). As these gene groups are stringently filtered and are not exhaustive, all of the genes not included in the groups described above were excluded from further analysis in our study.

ZGA timing groups were defined by the first timepoint that a gene is detected above a threshold of 5 TPM in the nascent dataset in any replicate. Genes that are never detected above 5 TPM in any replicate are labeled as Non-Zygotic.

### Statistical testing for association of gene groups

To quantify the significance of association between multiple gene lists, such as H3K4me3 dynamics groups, RNA dynamics groups and ZGA timing groups, Fisher’s exact test was used. The results are visualized in Balloon plots. Each dot represents the result of a Fisher’s exact test on a 2×2 contingency table. The table is created by taking one group of the first list of groups of interest and merging all other groups of the list into a comparison group. The same is repeated for the second list of groups of interest. The one-sided Fisher’s test is performed for positive association of the two gene lists of interest. The dot size and color show the FDR adjusted p-value to account for multiple testing.

### Gene ontology analysis

Gene ontology enrichment was calculated with the clusterProfiler package (v4.10.0) in R. The visualization of the similarities of the enriched GO terms (Benjamini-Hochberg adjusted p-value < 0.01) was done with the simplifyEnrichment package (v1.12.0).

### mRNA production

Four independent constructs: mouse Kdm5b (accession number NM_152895, aa1-770), its catalytic inactive (ci) mutant (H499A; aa1-770), Xenopus H3.3 (accession number NM_001098432) and its dominant-negative mutant (K4M) were sub-cloned into pCS2+ vector with Xenopus globin 5’ and 3’ UTRs followed by a polyadenylation signal-sequence of Simian Virus 40 (SV40), a C-terminus 3xHA tag and a C-terminus NLS-tag^13^. Additionally, the AID sequence from the pRN3P_AID-Dam-only construct (a kind gift by Maria-Elena Torres-Padilla; Addgene plasmid #136065)^61^ was subcloned into each of the four plasmids between the protein sequence and 3x HA tag using Gibson assembly (Table 1). For auxin-induced protein degradation, the pRN3P-TIR1-3×Myc construct (a kind gift from Maria-Elena Torres-Padilla; Addgene plasmid #119766)^61^ was used. mRNA for all five Kpn1-linearized constructs was synthesized from *in vitro* using the RiboMAX™ Large Scale SP6 RNA polymerase kit (Promega, #P1280) or the mMESSAGE mMACHINE T3 transcription kit (Thermo Fisher Scientific, #AM1348).

**Table 1.**
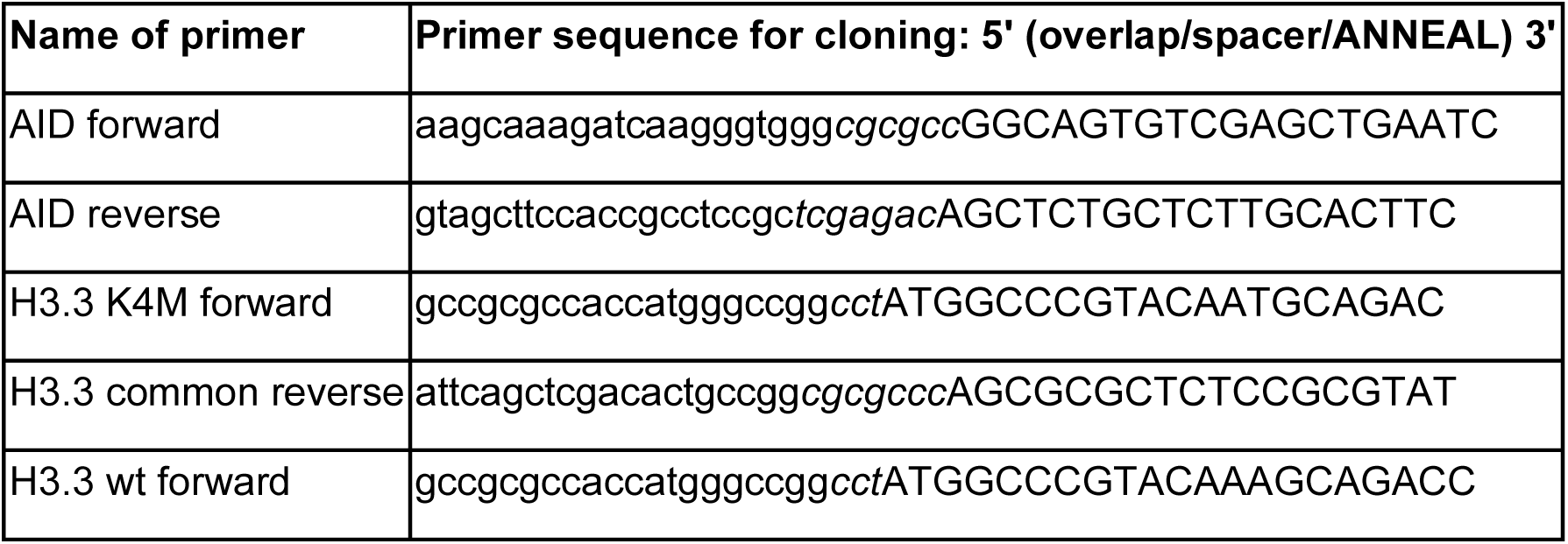

### Auxin experimental setup

For the “treated” condition, embryos were co-injected at the 1-cell stage with 11ng Kdm5b^wt^, 345pg H3.3^K4M^ and 5ng TIR1 mRNA. “Control” condition embryos were co-injected at the 1-cell stage with 4ng Kdm5b^ci^, 345pg H3.3^wt^ and 5ng TIR1 mRNA. mRNA amounts were determined on the basis of equal expression levels of the overexpressed proteins in pre-ZGA stage embryos. For “window” depletion treatment, embryos were grown until the pre-ZGA stage (4.5hpf/256-cell stage) and transferred to an auxin solution (500μM auxin in 0.1 × MMR) to develop further. “Persistent” depletion treatment embryos were grown in 0.1x MMR throughout development. To validate protein expression, pre-ZGA embryos were collected for Western blotting. To validate auxin-induced protein degradation, embryos were collected at the mid-ZGA stage (6.5hpf/4k-cell stage) and collected for Western blot and immunofluorescence. To assess the extent of H3K4me3 depletion, embryos were collected either at pre-ZGA stage for nuclear isolation and Western blot, or at NF12 for Western blot.

### Morpholino-mediated knockdown experimental setup

Custom antisense morpholinos (asMOs) targeting the translation start sites of both L and S homeologs of *Xenopus laevis kmt2b* and *cxxc1* were designed with the help of Gene Tools, LLC. The recommended standard control morpholino from Gene Tools LLC was used as a control. The morpholino sequences are shown in Table 2.

**Table 2.**
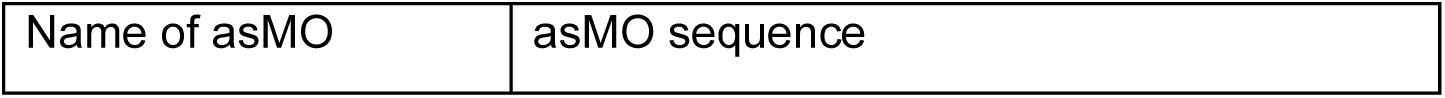
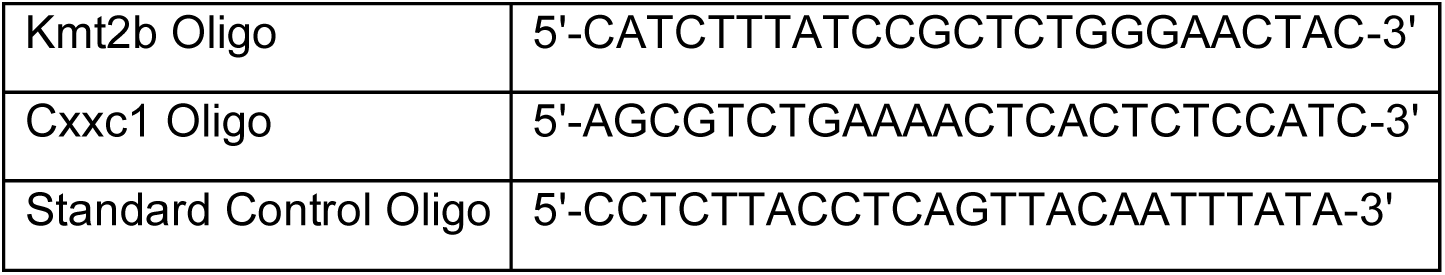

**Table 3.** Key resources used in this study.

| Reagent or resource | Source | Identifier |
| --- | --- | --- |
| **Primary antibodies** |  |  |
| H3K4me3 | Abcam | ab8580 |
| H3K4me1 | Abcam | ab8859 |
| H3K9ac | Millipore | #6-942 |
| H3K27ac | Cell Signalling Technologies | #8173 |
| H3K36me3 | Abcam | ab9050 |
| H3K79me3 | Diagenode | C15410068 |
| H4 | Abcam | ab31830 |
| Cfp1 | Biomol | A303-161A |
| Kmt2b/MLL2 | Abcam | ab104444 |
| **Secondary antibodies** |  |  |
| IRDye 680LT (donkey anti-mouse) | LICOR | 926-68022 |
| IRDye 800CW (donkey anti-rabbit) | LICOR | 926-32213 |

| **Kits** |  |  |
| --- | --- | --- |
| Methylated DNA Capture kit | Diagenode | C02020010 |
| NEBNext Ultra II Directional RNA Library Prep Kit for Illumina | NEB | E7760L |
| NEBNext Multiplex Oligos for Illumina Set 1/2/3 (96 Unique Dual Index Primer Pairs) | NEB | E6440S/E6442S/E6444S |
| **Software** | **Version** |  |
| GraphPad Prism | 10 |  |
| IGV | 2.13.0 |  |
| Fiji ImageJ | 2.14.0/1.54f |  |
| ImageStudio | 5.5.4 |  |
| SnapGene Viewer | 7.1.1 |  |

1-cell stage embryos were injected with 9.2nl of 1mM morpholino solution and developed in 0.5x MMR solution for 2 hours before moving to 0.1x MMR. Embryos were grown at 18°C until the desired stage. For analysis of developmental phenotypes, embryos were imaged and counted at periodic intervals with daily replacement of 0.1x MMR.

### RNA-seq experimental setup

For RNA-sequencing experiments, independent experiments were performed on two different frogs, serving as two biological replicates. Within each biological replicate, three sets of three embryos each were collected per time point, corresponding to three replicates within each independent experiment. Embryos were collected as described at hourly intervals after the pre-ZGA stage (i.e. 4.5hpf/256-cell stage): at 5.5hpf, 6.5hpf and 7.5hpf. All liquid was removed from the tube and samples were snap-frozen on dry ice and then stored at −80°C until further processing. Gene expression was analyzed for morpholino knockdown experiments and H3K4me3 window-depletion experiments.

### Total RNA extraction and ribosomal depletion

Three embryos were collected per sample and stored at −80°C. Total RNA was isolated using the RNeasy kit (Qiagen, #74106) according to the manufacturer’s instructions. First, embryos were lysed in the RLT buffer by vortexing for 15 minutes at 4°C. DNase digestion was performed according to the manufacturer’s instructions using RNase-free DNase (Qiagen, #79254). RNA was eluted in 40µl nuclease-free H2O. Concentration was measured using Nanodrop. Per sample, 500ng of total RNA was used for ribosomal RNA depletion. Ribosomal RNA was depleted using custom-made oligomer mixes for *Xenopus laevis* rRNA as described previously^65^ with minor modifications. Briefly, RNA was hybridized with 1 µL each of probe mix 1 and probe mix 2 in 20 µL hybridization buffer (100mM Tris–HCl pH 7.4, 200mM NaCl, 10mM DTT) by incubating at 95°C for 2 minutes. Hybridized samples were treated with 10 units of RNase H (NEB, #M0523) at 65°C for 5 minutes, followed by 2 units of RQ1 RNase-free DNase (Promega, #M6101) at 37°C for 30 minutes.

### RNA-seq library preparation and sequencing

Sequencing libraries were generated using the NEBNext Ultra II Directional RNA Library Preparation Kit for Illumina (NEB, #E7760) as per the manufacturer’s instructions using 12-13 PCR amplification cycles. The quality of cDNA libraries was assessed using the Agilent High Sensitivity D5000 ScreenTape System (Agilent, #5067-5592) in the Agilent 4150 TapeStation System or using the Agilent DNF-474 HS NGS Fragment Kit (Agilent, DNF-474-1000) in the Fragment Analyzer System (Agilent). Libraries were multiplexed and sequenced was performed as paired-end 50 bp reads on the Illumina NovaSeqX+ by the Helmholtz Core Facility Genomics (CF-GEN).

### RNA-seq data processing

Paired sequencing reads were processed using Kallisto (v0.48) for pseudoalignment and quantification of transcript abundance. Transcript and annotation files were downloaded from Xenbase (v10.1) for Xenopus laevis (transcripts and annotation). After quantification, transcript-level abundances were imported using ‘tximport’ (v1.32.0) and converted to a ‘SummarizedExperiment’ (v1.34.0) object in R (v4.3.2). Datasets of two independent batches were subjected to differential expression analysis on each of their three respective replicates performed with DESeq2 (v1.42.1). Reported p-values are FDR adjusted for multiple testing. Transcript abundances were normalized to transcripts per million (TPMs) and log2-transformedwith a pseudo count of 1 for further analysis.

## Data and Code availability

Original data produced in the experiments has been deposited on GEO. The MBD-seq data is available under the accession number GSE286525. The CATaDa data is available under the accession number GSE286524. The RNA-seq data from the auxin experiment is available under the accession number GSE286887. The RNA-seq data from the morpholino treatments is available under the accession number GSE286886. The mass spectrometry proteomics data have been deposited to the ProteomeXchange Consortium via the PRIDE^66^ partner repository with the dataset identifier PXD058937. Publicly available data were obtained from GEO with the following accession numbers: GSE125982, GSE75164, GSE76059, GSE201835. All original code can be accessed at https://github.com/ScialdoneLab/h3k4me3_maintenance. The preprocessed data for the analysis can be downloaded from Zenodo (https://doi.org/10.5281/zenodo.14648394). Any additional information required to reanalyze the data reported in this paper is available from the lead contact upon request.

## Acknowledgements

M.S.O. and E.H. were supported by the HO 6864/2-1 project DFG grant; CRC1064 Chromatin Dynamics Project-ID 213249687, Recognition Award of the Helmholtz Association; Project grant of the MRC MR/P00479/ S.H. by ERC starting grant 852798. Work in the labs of E.H. and A.S. was funded by the Helmholtz Association. A.E. received support by the Helmholtz Association awarded to Maria-Elena Torres-Padilla. M.S. was supported by the “Joachim Herz Stiftung” fellowship and the “Munich School for Data Science—MUDS”. JvdA is supported by a Wellcome Clinical Research Career Development Fellowship (219615/Z/19/Z), Wellcome Discovery Award (226653/Z/22/Z), a UKRI BBSRC Grant (BB/X00256X/1) and Medical Research Council (MC_UU_00028/8). The authors are grateful to the members of the CAM LMU Munich and G. Eckstein and I. de la Rosa Velazquez from the Helmholtz Genomics Core Facility (CF-GEN) for technical support. We thank all present and past lab and institute members, especially Nemanja Vasovic and Ana Janeva, as well as Ralph Rupp and Kikue Tachibana for reagents, their critical input and support. For the purpose of open access, the author has applied a Creative Commons Attribution (CC BY) license to any Author Accepted Manuscript version arising from this submission.

## Author contributions

M.S.O performed experimental work with assistance from M.M. I.F. and A.I. performed and analyzed the mass spectrometry experiments, A.H.M and J.v.d.A performed and analyzed the CATaDa experiments. M.S., T.S. and M.S.O. performed computational analyses with input from A.S. and E.H. The manuscript was written by M.S.O. and E.H. with input from A.S. and M.S., and E.H. designed and supervised the study.

## SUPPLEMENTARY INFORMATION

**Figure S1.**
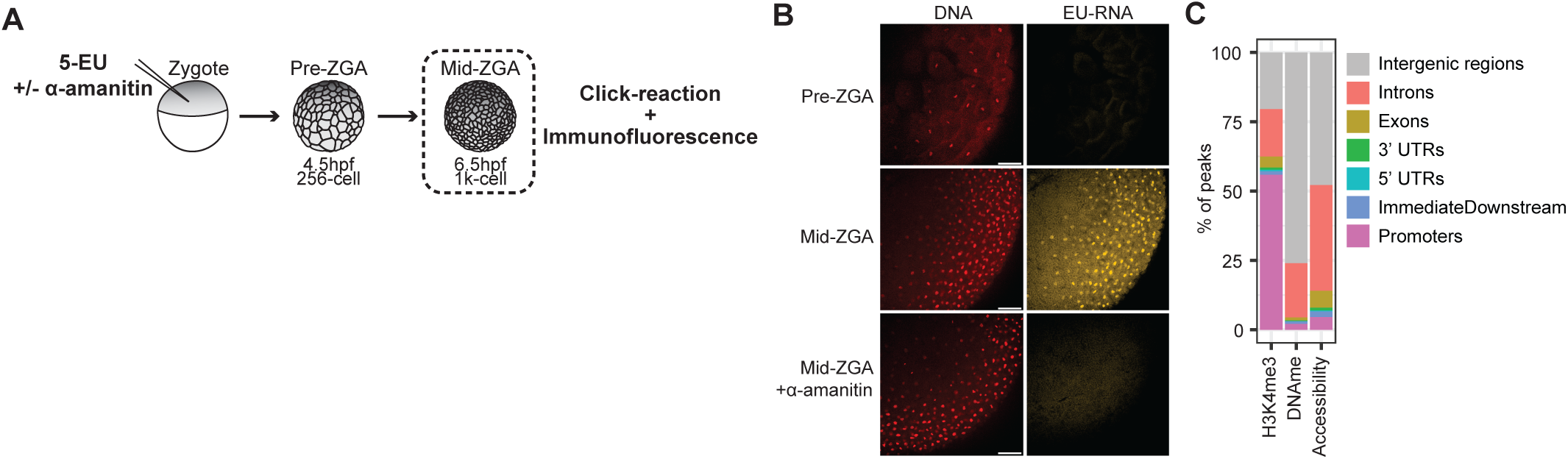
Profiling histone modifications and epigenetic factors present on pre-ZGA embryo chromatin. (A) Illustration of 5-EU labeling of nascent RNA in early embryos for immunofluorescence. (B) Representative images demonstrating inhibition of transcription ⍺-amanitin-treated embryos. Nascent transcripts are labeled using 5-EU (yellow) and DNA is labeled using Sir-DNA (red). Scale bars: 100µm. (C) Genomic distribution of called peaks for H3K4me3 ChIP sequencing, MBD sequencing and CATaDa on pre-ZGA embryos (4.5hpf).

**Figure S2.**
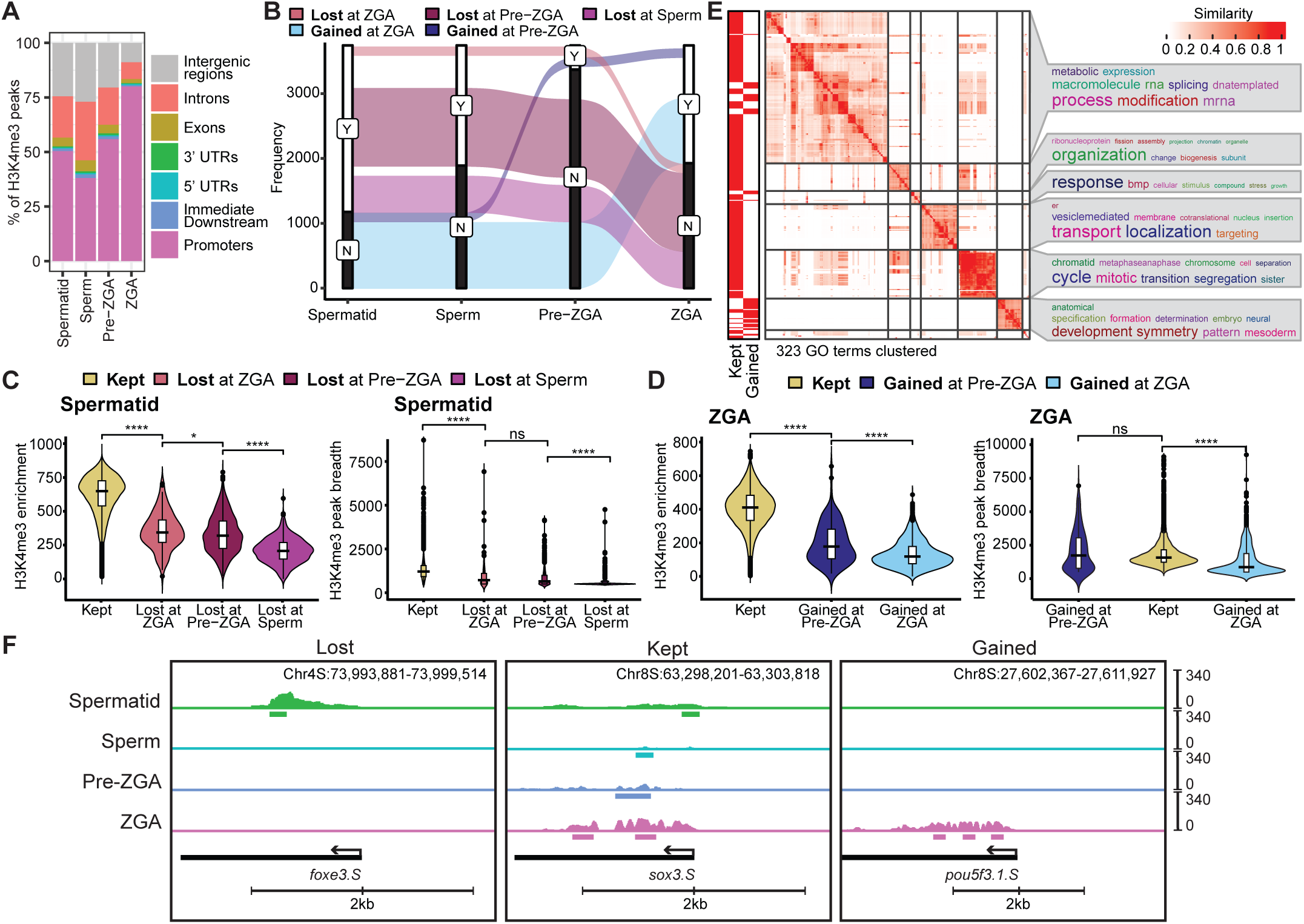
H3K4me3 peak intensity closely correlates with the duration of H3K4me3 maintenance. (A) Genomic distribution of H3K4me3 peaks at all time points. (B) Alluvial plot illustrating dynamics of promoter H3K4me3 for the GAINED and LOST groups. (C) Distribution of H3K4me3 promoter enrichment (left) and H3K4me3 peak breadth (bp) (right) at promoters for groups in spermatid Statistical test: one-sided Wilcoxon rank-sum test; p-values: (****)<=0.0001, (***)<=0.001, (**) <= 0.01, (*)<=0.05; n.s. are p-values > 0.05. (D) Distribution of H3K4me3 promoter enrichment (left) and H3K4me3 peak breadth (bp) (right) at promoters for groups in post-ZGA. (E) GO analysis for the KEPT and GAINED H3K4me3 dynamics groups that clusters significantly enriched GO terms (adj. p-value < 0.01) based on their similarity. For the LOST group, no GO term is significantly enriched. (F) IGV tracks of genes representing LOST, KEPT and GAINED H3K4me3 dynamics groups.

**Figure S3.**
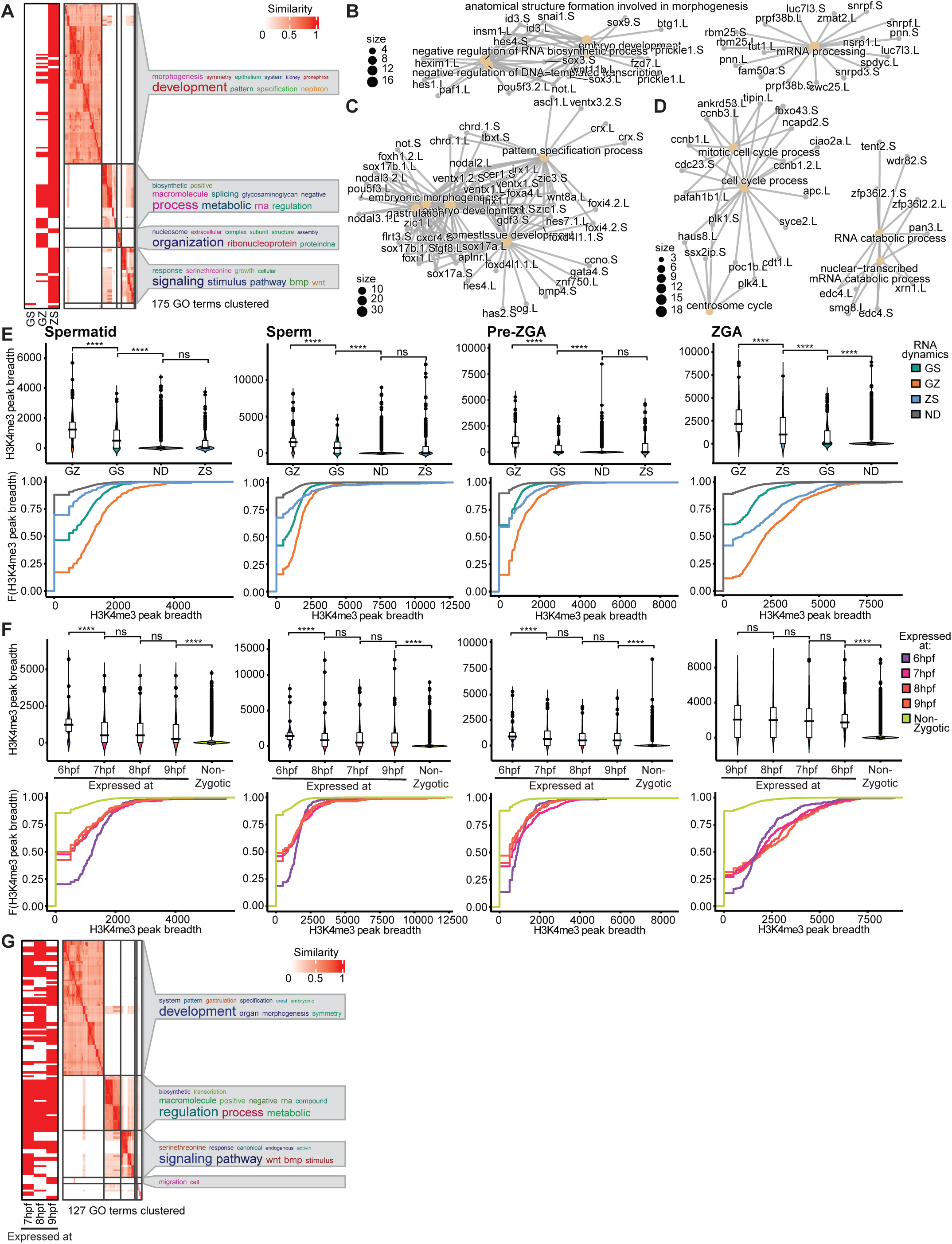
H3K4me3 is maintained independently of transcription between two transcriptionally active time points. (A) GO analysis for the different RNA dynamics groups that clusters significantly enriched GO terms (adj. p-value < 0.01) based on their similarity. (B-D) Networks showing enriched GO terms, their relation and associated genes for *(B) Gamete-Zygotic (GZ) genes, (C) Zygotic-Specific (ZS) genes and (D) Gamete-Specific (GS) genes.* (E) Distributions (Top) and cumulative distributions (ECDF plots) (Bottom) of H3K4me3 peak breadth (bp) at promoters for the RNA dynamics groups at each time point - spermatid, sperm, pre-ZGA, post-ZGA respectively. Statistical test: one-sided Wilcoxon rank-sum test; p-values: (****)<=0.0001, (***)<=0.001, (**) <= 0.01, (*)<=0.05; n.s. are p-values > 0.05. (F) Distributions (Top) and cumulative distributions (ECDF plots) (Bottom) of H3K4me3 peak breadth (bp) at promoters for the ZGA timing groups at each time point - spermatid, sperm, pre-ZGA, post-ZGA respectively. (G) GO analysis for the different ZGA timing groups that clusters significantly enriched GO terms (adj. p-value < 0.01) based on their similarity. For the 6hpf group, no GO term is significantly enriched.

**Figure S4.**
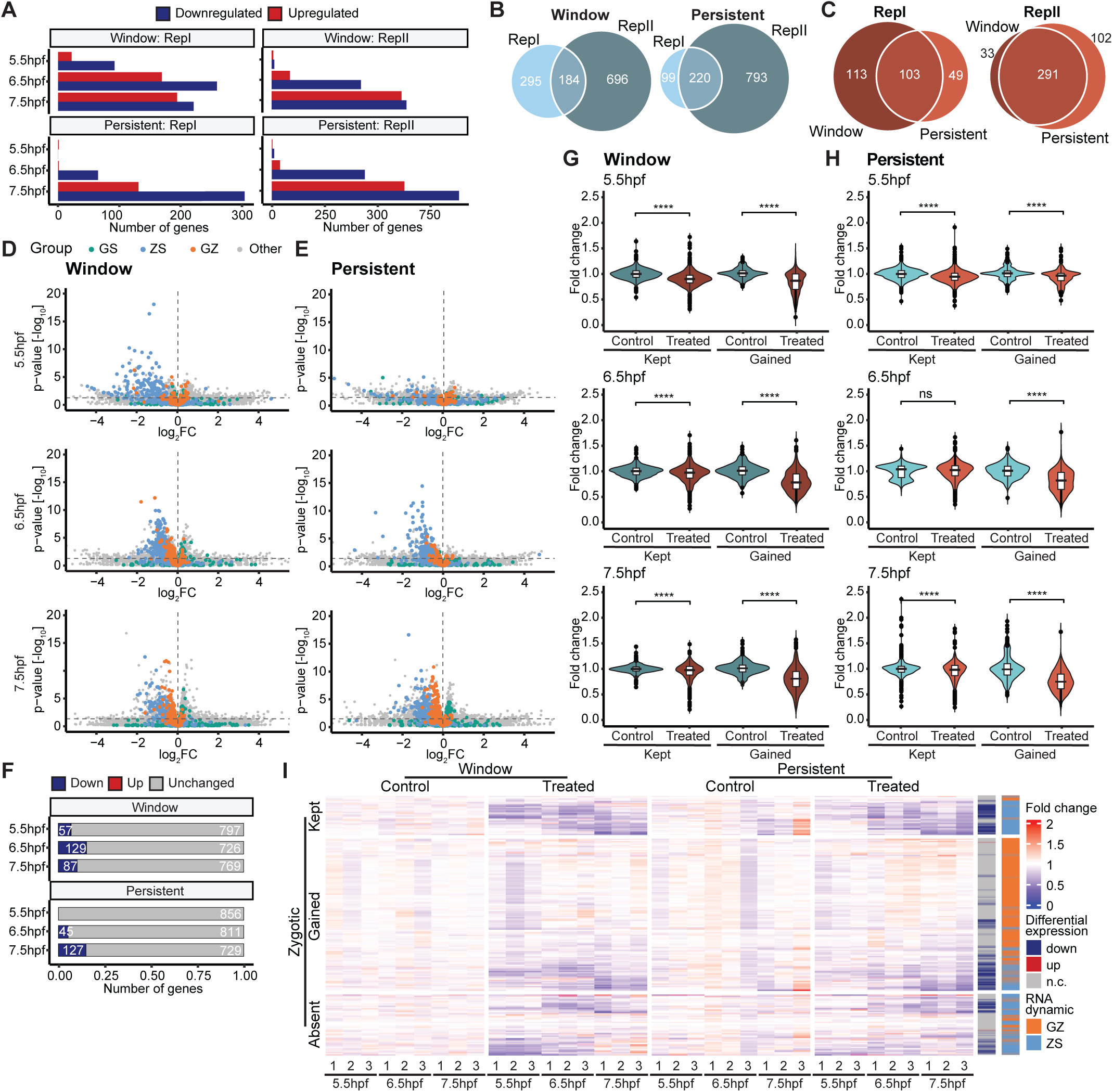
Maintenance of H3K4me3 during early embryonic cell divisions is required for proper ZGA. (A) Count of differentially expressed genes at 5.5hpf, 6.5hpf and 7.5hpf time points in window condition compared to the respective control samples for both biological replicates (adj. p-value < 0.05). (B) Overlap of downregulated genes at any timepoint in window and persistent depletion conditions compared to controls between biological replicates. (C) Overlap of downregulated genes at any timepoint between window and persistent depletion conditions compared to control in both replicates. (D-E) Volcano plots displaying expression fold-change and adj. p-value (cutoff: 0.05) in treated condition compared to control condition at each timepoint colored by RNA dynamics groups for (D) window depletion and (E) persistent depletion conditions. (F) Number of differentially expressed and unchanged zygotic genes at 5.5hpf, 6.5hpf and 7.5hpf time points in window treatment and persistent treatment compared to control embryos in biological replicate 2 (adj. p-value < 0.05). Zygotic genes = Gamete-Zygotic (GZ) + Zygotic-Specific (ZS). (G-H) Fold-change of gene expression levels of all zygotic genes in each group calculated over mean TPM of respective control technical replicates for (G: Window, H: Persistent) at the three time points in biological replicate 2. Statistical test: one-sided Wilcoxon rank-sum test; p-values: (****)<=0.0001, (***)<=0.001, (**) <= 0.01, (*)<=0.05; n.s. are p-values > 0.05. (I) Expression of zygotic genes in 3 technical replicates of every sample, clustered by H3K4me3 dynamics group (KEPT, GAINED and ABSENT) for biological replicate 2. Fold-change of every gene is calculated over mean TPM of three technical replicates of the control condition of each respective time point. Genes are annotated by differential gene expression, and RNA dynamics group.

**Figure S5.**
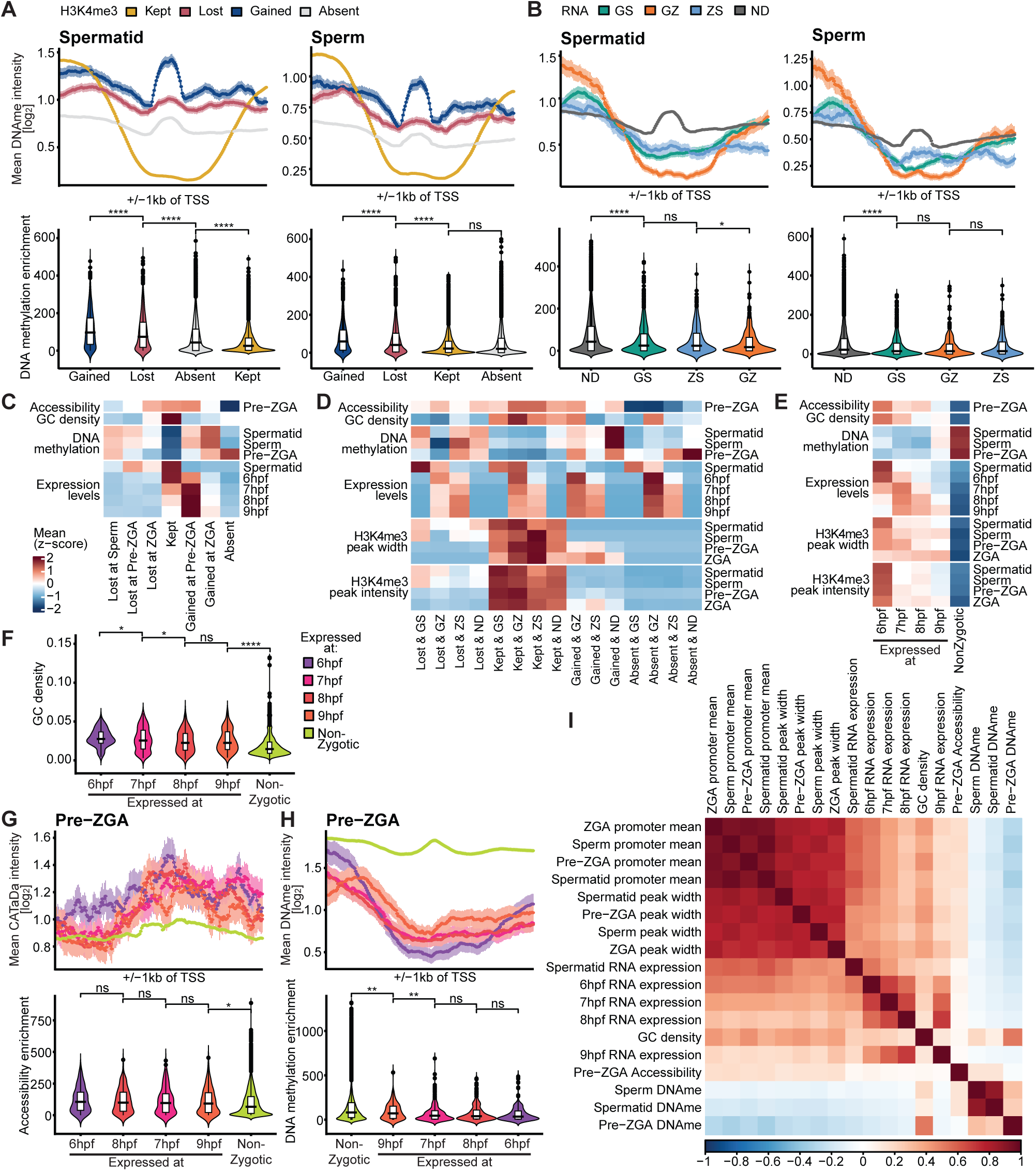
H3K4me3 maintenance around promoters correlates with increased chromatin accessibility, high GC content and DNA hypomethylation. (A-B) (Top) DNA methylation around TSS (+/−1kb) at spermatid and sperm stages. (Bottom) DNA methylation promoter enrichment in each group for spermatid and sperm stages. (A: H3K4me3 dynamics groups; B: RNA dynamics groups). Statistical test: one-sided Wilcoxon rank-sum test; p-values: (****)<=0.0001, (***)<=0.001, (**) <= 0.01, (*)<=0.05; n.s. are p-values > 0.05. (C) Heatmap showing the z-scored mean of accessibility, CG density and DNA methylation at promoters and expression levels for available timepoints for the different H3K4me3 dynamics groups. (D) Heatmap showing the z-scored mean of accessibility, CG density, DNA methylation, H3K4me3 enrichment and peak width at promoters and expression levels for available timepoints for the intersections of major H3K4me3 dynamics groups with RNA dynamics groups. (E) Heatmap showing the z-scored mean of accessibility, CG density, DNA methylation, H3K4me3 enrichment and peak width at promoters and expression levels for available timepoints for the different ZGA timing groups. (F) Comparison of promoter CG density for ZGA timing groups. (G) Accessibility measured by DamID intensity around TSS (+/−1kb) for ZGA timing groups at pre-ZGA stage. (Bottom) accessibility promoter enrichment in each group for pre-ZGA stage. (H) DNA methylation around TSS (+/−1kb) for ZGA timing groups at pre-ZGA stage. (Bottom) DNA methylation promoter enrichment in each group for pre-ZGA stage. (I) Pairwise linear correlation computed between all presented data modalities.

**Figure S6.**
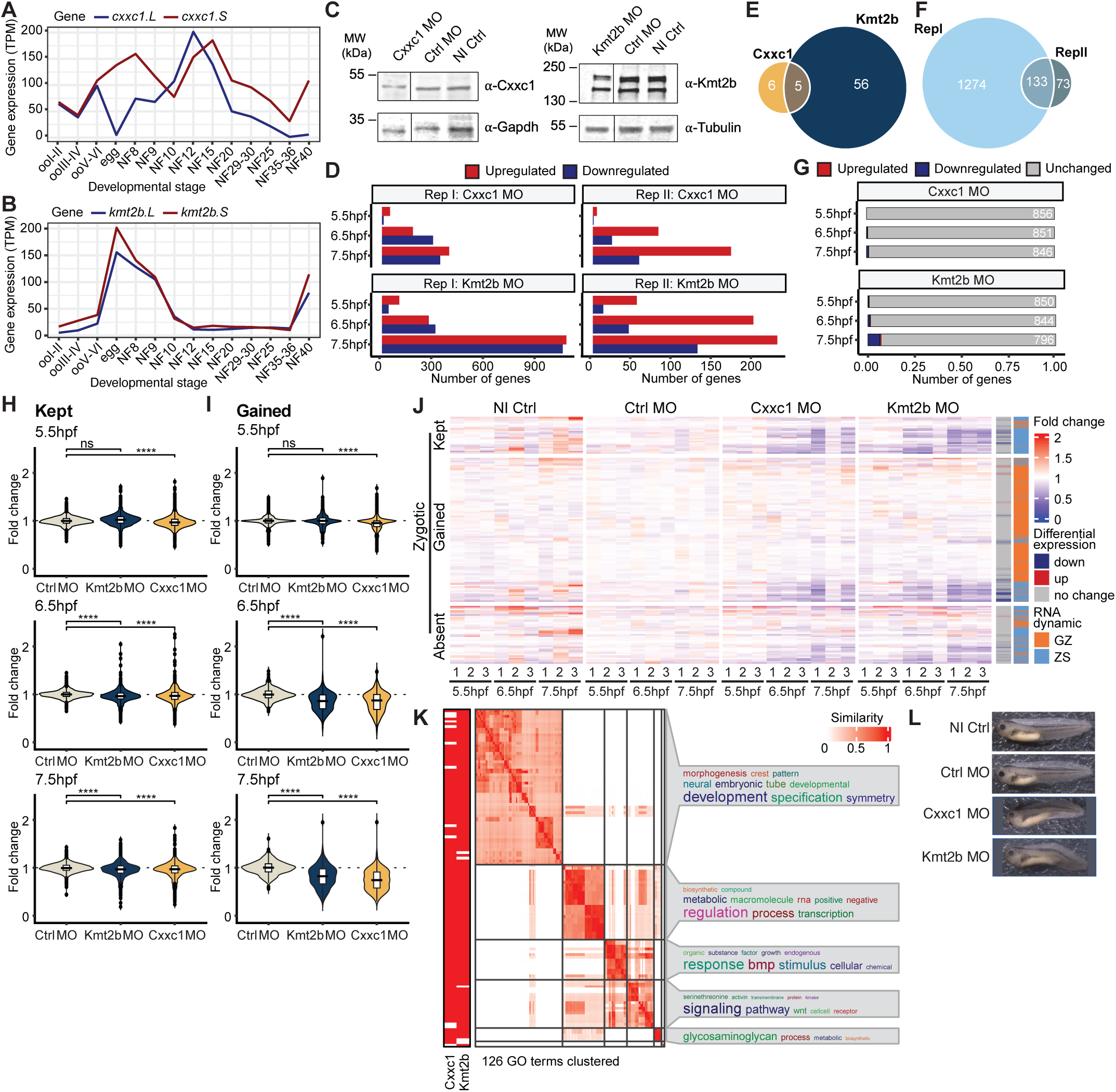
Cxxc1 and Kmt2b are required for proper ZGA and embryonic development. (A) Gene expression levels of *cxxc1.L* and *cxxc1.S* at stages of development (Session et al., 2016). (B) Gene expression levels of *kmt2b.L* and *kmt2b.S* at stages of development (Session et al., 2016). (C) Western blots showing Cxxc1 and Kmt2b protein levels in Cxxc1 and Kmt2b knockdown embryos. (D) Count of differentially expressed genes at 5.5hpf, 6.5hpf and 7.5hpf time points in Cxxc1 and Kmt2b morpholino-injected conditions compared to control morpholino embryos (adj. p-value < 0.05). (E) Overlap of downregulated zygotic genes at any timepoint between Cxxc1 and Kmt2b morpholino conditions compared to control morpholino. (F) Overlap of downregulated genes compared to control morpholino at any timepoint between both biological replicates. (G) Number of differentially expressed and unchanged zygotic genes at 5.5hpf, 6.5hpf and 7.5hpf time points in Cxxc1 and Kmt2b morpholino-injected conditions compared to control morpholino embryos in biological replicate 2 (adj. p-value < 0.05). *Zygotic genes = Gamete-Zygotic (GZ) + Zygotic-Specific (ZS).* (H-I) Fold-change of gene expression levels of all zygotic genes in each group calculated over mean TPM of control morpholino technical replicates for (H) KEPT and (I) GAINED groups at the three time points in biological replicate 2. Statistical test: one-sided Wilcoxon rank-sum test; p-values: (****)<=0.0001, (***)<=0.001, (**) <= 0.01, (*)<=0.05; n.s. are p-values > 0.05. (J) Expression of zygotic genes in 3 technical replicates of every sample, clustered by H3K4me3 dynamics group (KEPT, GAINED and ABSENT) for biological replicate 2. Fold-change of every gene is calculated over mean TPM of three technical replicates of control morpholino of the respective time point. Genes are annotated by differential gene expression, and RNA dynamics group. (K) Gene ontology analysis of downregulated zygotic genes from Fig.4G. (L) Representative phenotype images of embryos for each condition at NF41. n=2 experiments, N>35 per condition.

**Table S1.** Number of genes overlapping between each H3K4me3 dynamics group and RNA dynamics group.

|  |  | RNA dynamics groups |  |  |  |
| --- | --- | --- | --- | --- | --- |
|  |  | Gamete-Specific | Gamete+Zygotic | Zygotic-Specific | Not-detected |
| H3K4me3 dynamics groups | LOST | 122 | 1 | 0 | 585 |
|  | KEPT | 320 | 377 | 96 | 1112 |
|  | GAINED | 0 | 16 | 105 | 382 |
|  | ABSENT | 329 | 49 | 140 | 12100 |

